# Rhythms of Transcription in Field-Grown Sugarcane Are Highly Organ Specific

**DOI:** 10.1101/607002

**Authors:** Luíza Lane de Barros Dantas, Felipe Marcelo Almeida-Jesus, Natalia Oliveira de Lima, Cícero Alves-Lima, Milton Yutaka Nishiyama, Monalisa Sampaio Carneiro, Glaucia Mendes Souza, Carlos Takeshi Hotta

**Affiliations:** Departamento de Bioquímica, Instituto de Química, Universidade de São Paulo, São Paulo, SP, 05508-000, Brazil; Max Planck Institute for Molecular Plant Physiology, Potsdam-Golm, 14476, Germany; Laboratório Especial de Toxicologia Aplicada, Instituto Butantan, São Paulo, SP, 05503-900, Brazil; Departamento de Biotecnologia e Produção Vegetal e Animal, Centro de Ciências Agrárias, Universidade Federal de São Carlos, São Carlos, SP, 13600-970, Brazil

## Abstract

We investigated whether different specialized organs in field-grown sugarcane follow the same temporal rhythms in transcription. We assayed the transcriptomes of three organs during the day: leaf, a source organ; internodes 1 and 2, sink organs focused on cell division and elongation; and internode 5, a sink organ focused on sucrose storage. The leaf had twice as many rhythmic transcripts (>68%) as internodes, and the rhythmic transcriptomes of the two internodes were more similar to each other than to those of the leaves. More transcripts were rhythmic under field conditions than under circadian conditions and most of their peaks were during the day. Among the transcripts that were considered expressed in all three organs, only 7.4% showed the same rhythmic time course pattern. The central oscillators of these three organs — the networks that generate circadian rhythms — had similar dynamics with different amplitudes. The differences between the rhythmic transcriptomes in circadian conditions and field conditions highlight the importance of field experiments to understand the plant circadian clock *in natura*. The highly specialized nature of the rhythmic transcriptomes in sugarcane organs probably arises from amplitude differences in tissue-specific circadian clocks and different sensitivities to environmental cues.

**One sentence summary:** The rhythmic transcriptome of field-grown sugarcane is highly organ-specific.

## Introduction

The circadian clock is an endogenous signaling network that allows organisms to adapt to rhythmically changing environments. Plants with a circadian clock synchronized with environmental rhythms accumulate more biomass and have better fitness than plants with defective or no circadian clocks^1,2^. In crops, changes in the circadian clock have been indirectly selected through traditional breeding to change photoperiodic responses, such as the transition to flowering. For example, the circadian clocks of European tomatoes have longer periods than those of native American tomatoes, as such periods allow these crops to adapt better to the long summer days occurring at the high latitudes of much of Europe^3^. Similarly, some genotypes of *Hordeum vulgare* L. (barley) and *Triticum aestivum* L. (wheat) carry mutations in their circadian clock genes that reduce flowering induced by photoperiodic triggers, allowing cultivation in higher latitudes in Europe^4,5^.

The circadian clock is conceptually divided into three associated parts: the *Input Pathways,* the *Central Oscillator,* and the *Output Pathways.* The *Input Pathways* detect entraining cues that keep the circadian clock continuously synchronized to the environment. In plants, these cues include light, temperature, and sugar levels^6–8^. The *Central Oscillator* is a series of interlocking transcriptional-translational feedback loops that can generate 24-h rhythms independently of the environment. In *Arabidopsis thaliana* (L.) Heynh. (Arabidopsis), one loop, called the morning loop, starts with the light induction of *CIRCADIAN CLOCK ASSOCIATED1 (CCA1)* and *LATE ELONGATED HYPOCOTYL (LHY)* at dawn. Next, *PSEUDO-RESPONSE REGULATOR7 (PRR7)* and *PRR9* are activated by CCA1 and LHY. In turn, *CCA1* and *LHY* are repressed by PRR7 and PRR9. In the core loop, CCA1 and LHY are repressed, and this represses *TIME FOR CAB EXPRESSION1 (TOC1),* also known as *PRR1.* During the night, TOC1 forms an interaction known as the evening loop with the EVENING COMPLEX (EC). The EC is a protein complex formed by EARLY FLOWERING3 (ELF3), ELF4, and LUX ARRHYTHMO (LUX) that also inhibits the expression of *PRR7* and *PRR9* the next morning. Other essential components of the oscillator include GIGANTEA (GI), REVEILLE8 (RVE8), and CCA1 HIKING EXPEDITION (CHE)^8–11^. The *Output Pathways* transduce the temporal information generated by the interaction between the *Central Oscillator* and the *Input Pathways* to a plethora of biochemical pathways. The circadian clock thus has a broad impact throughout the plant, regulating processes such as photosynthesis, cell elongation, stomata opening, and flowering^12^.

Even though the plant circadian clock is highly conserved, there are a few differences between the circadian clocks of Arabidopsis and grasses (Poales). For instance, there is only copy of the paralogs *CCA1/LHY,* usually assigned as *LHY*^13^ The grass PRRs consist of *TOC1, PRR37, PRR73, PRR59,* and *PRR95,* and it is not clear whether they have the same functions as their Arabidopsis counterparts, even though they are capable of complementing Arabidopsis mutations^13,14^. In sugarcane, a highly polyploid crop that accumulates sucrose in the culm, the circadian clock has high-amplitude rhythms and regulates a large proportion of the leaf transcriptome (>30%)^15,16^.

Most research to date on plant circadian rhythms has been done in controlled conditions, inside a growth room or growth chamber. Under such circumstances, plants can be grown either under circadian conditions, in which they are kept under constant abiotic conditions as a means to separate endogenous rhythms from rhythms driven by the environment, or under diel conditions, in which they are subjected to abiotic rhythms such as light/dark and warm/cold. Abiotic changes in controlled conditions are usually stepwise, in contrast to the gradients found in natural or field conditions, which can lead to significant changes in plant physiology^17–19^. For example, different patterns of metabolite rhythms are observed if plants are grown under white fluorescent tubes, light-emitting diodes that simulate the sunlight spectrum, or naturally illuminated greenhouse^18^. In another study, the period and phase of the circadian clock affected shoot and rosette branch numbers in multiple Arabidopsis mutants in natural, but not controlled, conditions^20^. Finally, the rice mutant *osgi,* which has a late-flowering phenotype in controlled conditions, flowered at the same time as the wild type in field conditions^21^.

Only two plant species have had their rhythmic transcripts identified in field conditions: rice and pineapple^21–24^. However, these studies focused on the leaves. To better understand how the plant circadian clock regulates transcription under natural conditions in different organs, we measured transcription in three organs of field-grown sugarcane grown during the day. We harvested leaf +1 (L1), a source organ, and two sink organs: internodes 1 and 2 (I1), organs focused on cell division and cell elongation that includes the shoot apical meristem; and internode 5 (I5), an organ focused on sucrose accumulation. We describe in detail one cycle (24 h) with 14 time points, starting 2 h before dawn. This approach allowed us to obtain a better resolution to describe transcripts with fast dynamics. We found that the rhythmic transcripts of the L1, I1, and I5 are widely specialized and likely to respond differently to environmental cues.

## Results

### A significant proportion of the sugarcane transcriptome is rhythmic in diel conditions

We planted a field of commercial sugarcane (*Saccharum* hybrid SP80-3280) in autumn 2012 in Araras (Brazil, 22°18′41.0”S, 47°23′5.0″W). Nine months later (summer 2013), after a dry winter and spring (Fig. S1), we did a time course experiment in which the leaf +1 (L1), internodes 1 and 2 (I1), and internode 5 (I5) were harvested every 2 h for 26 h, starting 2 h before dawn. On the day of harvest, the stalks were 76 ± 0.16 cm, with 11 ± 2 internodes, and their sugar content was 12.0 ± 1.4°Bx (mean ± SD; *n* = 20). The temperature varied throughout the day from 17°C to 30°C, with the maximum occurring at 11 h after dawn (ZT11); the maximum light intensity was 2.67 MJ/m^2^ at ZT07, and dusk occurred 13.25 h after dawn (ZT13.25) (Fig. S1B and D).

RNA extracted from each organ was hybridized in 44k custom oligoarrays^15,25^. The data from the time course experiment generated 14,521 time series with 14 time points. After the selection of time points that had a signal above the background noise (Figure S3A), we had 12,501 transcripts considered to be expressed in at least one organ (Fig. 1). L1 had 9,822 expressed transcripts, 94.3% of them were also expressed in a previous circadian experiment^15^ (Fig. 1B). I1 had the highest number of expressed transcripts (12,053), followed by I5 (10,448). A total of 9,380 transcripts were expressed in all three organs (75.0%, Fig. 1E). I1 and I5 shared the most substantial proportion of the expressed transcripts (89.3%), and I1 had the most substantial proportion of unique expressed transcripts (7.5%).

**Figure 1 –.**
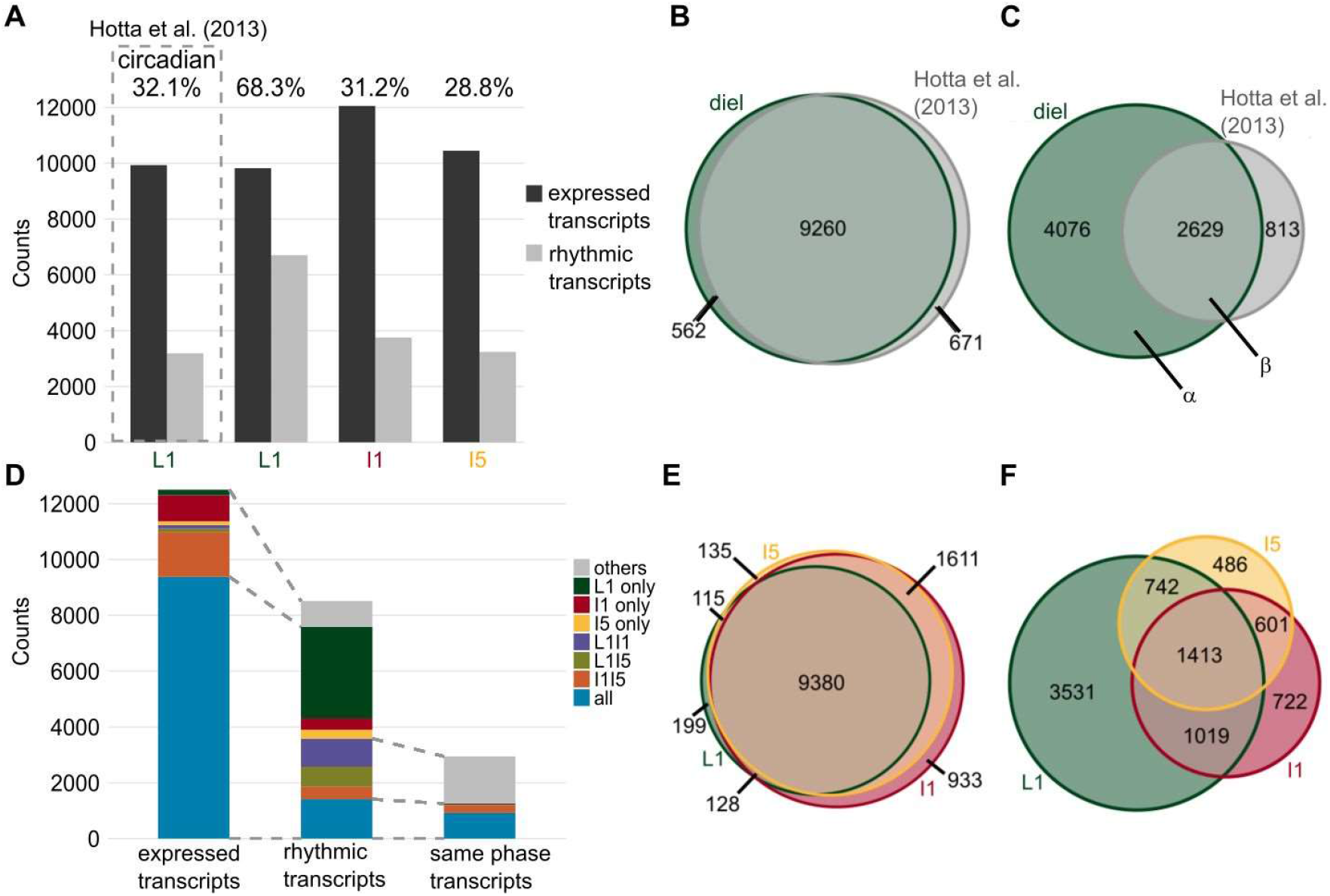
Different organs have specific sets of rhythmic transcripts in sugarcane. **(A)** The numbers of expressed and rhythmic transcripts detected in leaf +1 (L1), internodes 1 and 2 (I1), and internode 5 (I5) in field-grown (diel) conditions, and in leaf +1 in circadian conditions published in Hotta et al. (2013)^15^. **(B, C)** Euler diagrams of expressed transcripts **(B)** and rhythmic transcripts **(C)** in L1 in sugarcane in diel (green) and circadian (gray) conditions. **(D)** Number of expressed transcripts, rhythmic transcripts, and rhythmic transcripts with the same phase that were found specifically in L1, I1, or I5; in both L1 and I1 (L1I1, purple); in both L1 and I5 (L1I5, light green); in both I1 and I5 (I1I5, orange); and in all three organs (L1I1I5, blue). In the second bar, the gray area corresponds to rhythmic transcripts that are expressed in only one or two organs. In the third bar, the gray area corresponds to rhythmic transcripts in only two organs that have the same phase. The gray dashed lines show the associations among bars. **(E, F)** Euler diagram of expressed and rhythmic transcripts in L1, I1, and I5 in field-grown sugarcane in diel conditions.

We identified rhythmic transcripts by combining a weighted correlation network analysis (WGCNA) that grouped expressed transcripts in coexpression modules^26^ with JTK_CYCLE, which identified which of the modules contained rhythmic transcripts^27^ (Figure S3B). This method identified 6,705 rhythmic transcripts in L1 (68.3%), 3,755 in I1 (31.2%), and 3,242 in I5 (28.8%) (Fig. 1A and Fig. S6). As a comparison, 32.1% of the transcripts were rhythmic in L1 under circadian conditions^15^. The overlap between circadian transcripts and rhythmic transcripts in the field (in diel conditions) was 2,623, representing 76.4% of circadian transcripts and 60.1% of rhythmic transcripts (Fig. 1C).

### Different sets of transcripts are rhythmic in different sugarcane organs

Although most expressed transcripts were found in all three organs, only 1,413 of the expressed transcripts were rhythmic in all three organs (16.6%) (Fig. 1D, F). L1 had the largest proportion of unique rhythmic transcripts (41.5%), followed by I1 (8.5%) and then I5 (5.7%) (Fig. 1F). Transcripts that were expressed only in one organ were less likely to be rhythmic (60.3% for L1, 8.6% for I1, and 8.4% for I5) (Fig. 1H).

We estimated the phase of the transcripts by combining the phase calculated using JTK_CYCLE with a dendrogram of the representative time course of each module. Among the transcripts that were rhythmic in more than one organ, 27% had rhythms with phase differences >2 h (Fig. 1D). Overall, among the 12,501 unique expressed transcripts in the three organs, only 7.4% (923) showed rhythms with the same phase in the three organs. Most of the transcripts peaked during the day: this was true of 80.3% in L1, 90.4% in I1, and 96.3% in I5 (the photoperiod was 13.25 h, or 56.3% of a cycle) (Fig. 1G). In L1, 2,363 transcripts peaked between dawn (ZT00) and 2 h after dawn (ZT02) (35.2%), and 1,232 transcripts peaked at ZT12 (18.4%) (Fig. 2A). When we separated rhythmic L1 transcripts into those that were also rhythmic in circadian conditions (Fig. 2B, α) and those that were not (Fig. 2B, β), two different phase distributions could be observed (Fig. 2C). The α group had most transcripts peaking at ZT00-02 (39.1%), followed by ZT12 (14.0%), while the β group peaked at ZT12 (25.1%), followed by ZT02 (19.1%). In I1, 1,201 transcripts peaked at ZT0 (32.0%) and 716 peaked at ZT8 (19.1%). In I5, 1,373 transcripts peaked at ZT0 (42.4%) and 894 peaked at ZT8 (27.6%) (Fig. 2A).

**Figure 2 –.**
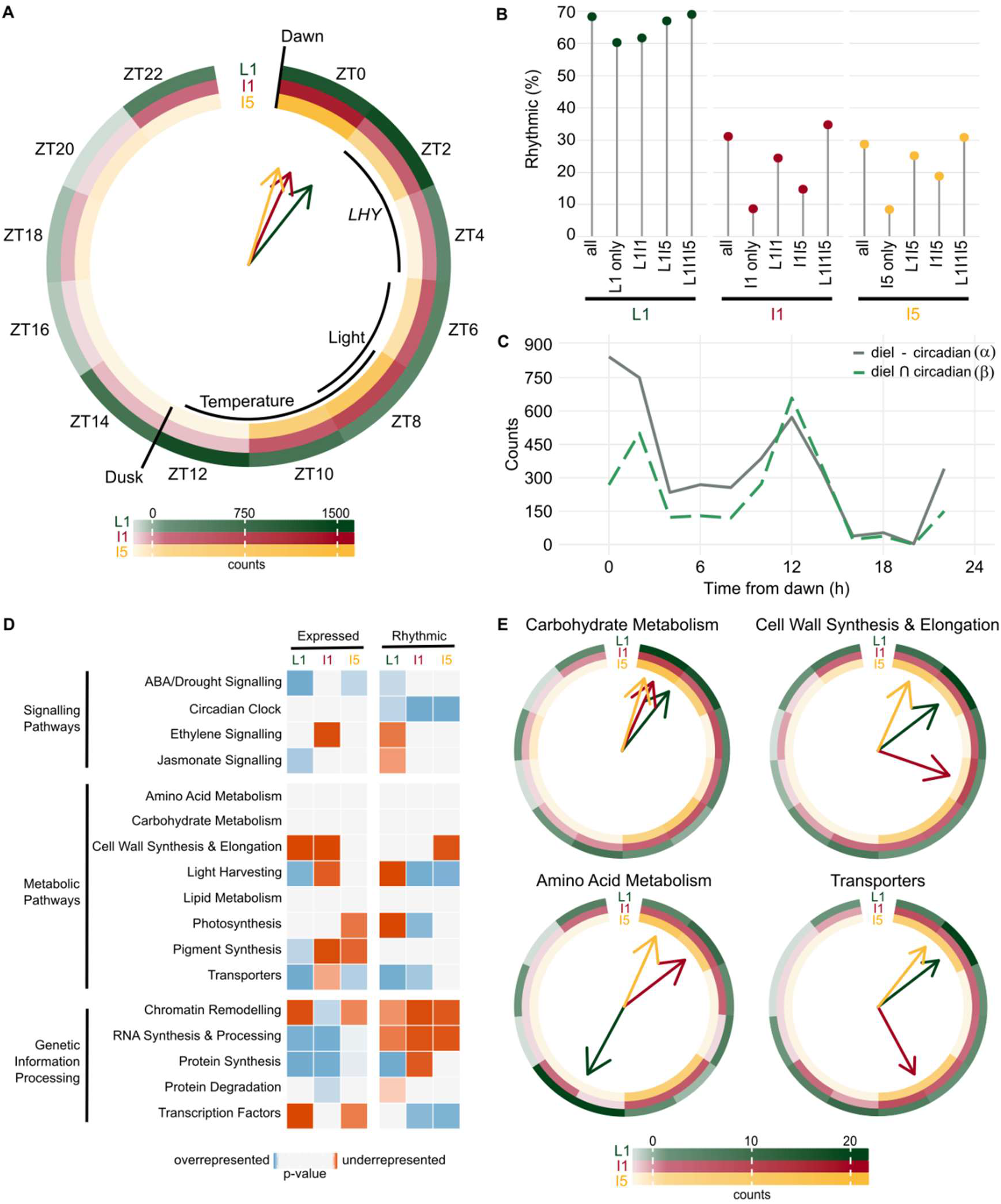
Transcripts have unique phases in different sugarcane organs. **(A)** Circular heatmap of the rhythmic transcript peak time (ZT0 = 0 h after dawn) distribution in leaf +1 (L1), internodes 1 and 2 (I1), and internode 5 (I5). The colored arrows show the times at which the most transcripts are found in each organ. The times of dawn, dusk, *LHY* transcription peak, maximum light intensity, and maximum temperatures are indicated by black arcs. **(B)** Proportions of transcripts that were rhythmic in L1, I1, and I5 among all expressed transcripts in each organ (All), among the transcripts expressed only in one organ (L1 only, I1 only, or I5 only), among the transcripts expressed in two organs (L1I1, L1I5, or I1I5), and among transcripts expressed in all three organs (L1I1I5). **(C)** Distribution of rhythmic transcript peak time in transcripts that were rhythmic in L1 but not in circadian conditions (α in Fig. 1C) and rhythmic transcripts in transcripts that were rhythmic in L1 and circadian conditions (β). **(D)** Heatmap of functional categories that are overrepresented (shades of blue) or underrepresented (shades of red) among the expressed and rhythmic transcripts of L1, I1, and I5. The P-value was calculated using a hypergeometric test. **(E)** Circular heatmap with the distribution of the peak times of rhythmic transcripts associated with the pathways *Carbohydrate Metabolism, Cell Wall Synthesis & Elongation, Amino Acid Metabolism,* and *Transporters.*

The majority of transcripts from L1 (65.8%) grown in diel conditions had the same phase (± 2 h) in leaves grown under circadian conditions (Fig. S7A). More transcripts showed a delayed peak (19.6%) rather than an advanced peak (13.9%) under diel conditions than under circadian conditions. When we compared L1 and I1 transcripts, 65.8% had the same phase, with the remainder divided roughly evenly between delayed and advanced phases (16.1% and 14.9%, respectively) (Fig. S7B). Similarly, 67.1% of the L1 transcripts had the same phase as I5, 14.2% had a delayed phase, and 14.8% had an advanced phase (Fig. S7C). The phases were most similar between I1 and I5 transcripts: 93.8% had the same phases, 2.8% a phase delay, and 3.1% a phase advance (Fig. S7D).

### Biochemical pathways have different rhythms in sugarcane organs

We used a hypergeometric test to detect if a pathway was over- or underrepresented by comparing the frequency of transcripts associated with a Biochemical Pathway among the expressed transcripts and all the unique transcripts in the oligoarray (Fig. 2D and Fig.). We used the same test comparing the frequency of transcripts associated with a Biochemical Pathway among the rhythmic transcripts and the expressed transcripts (Fig. 2D and Fig. S8). The transcript annotations were based on the SUCEST database annotation (http://sucest-fun.org).

Among expressed transcripts, each organ has a distinct profile. For example, L1 was the only organ that had the *Pigment Synthesis, Light Harvesting,* and *Jasmonate Signaling* pathways considered to be overrepresented. I1 had *Chromatin Remodeling* and *Protein Synthesis* pathways overrepresented and *Ethylene Signaling* underrepresented. *Transcription Factors* was underrepresented and *ABA/Drought Signaling* and *Transporters* were overrepresented in L1 and I5, but not in I1. I5 is the only organ in which *Cell Wall Synthesis & Elongation* was not underrepresented among the expressed transcripts (Fig. 2D). Among rhythmic transcripts, *Circadian Clock* was overrepresented, while *Chromatin Remodeling* and *RNA Synthesis & Processing* were underrepresented in all organs. *Protein Synthesis* was overrepresented in L1. *Transcription Factors* was overrepresented in I1 and I5, and *Transporters* was overrepresented among rhythmic transcripts in L1 and I1 (Fig. 2D).

When we analyzed transcripts associated with important pathways for sugarcane growth, we found further organ-specific patterns; these differences could be seen in both expressed and rhythmic transcripts, as well as the phase of the rhythmic transcripts (Fig. 2E and Fig. S9). Transcripts associated with *Carbohydrate Metabolism* tended to peak in the morning. Almost half (48.0%) of the transcripts had a peak at ZT00 in L1, while the majority peaked between ZT00 and ZT04 in both I1 (53.2%) and I5 (58%) (Fig. 2E). Amongst the individual transcripts, a putative ortholog of *SUCROSE SYNTHASE4 (SuSy4)* had a similar rhythmic pattern in all three organs. A putative ortholog of *SUCROSE-PHOSPHATE SYNTHASE II (SPSII)* was rhythmic only in L1, while a putative ortholog of a *CELL WALL INVERTASE (CWI)* exhibited a sharp peak at ZT04 in L1 but a very broad peak at ZT08 in I1 and I5 (Fig. S9I, M, and Q).

Transcripts associated with *Cell Wall Synthesis & Elongation* had a more diverse phase distribution: in L1, 55% had a peak between ZT00 and ZT04; in I1, 73.4% had a peak between ZT00 and ZT08; and in I5, 45.8% had a peak between ZT08 and ZT10 and 37.8% had one at ZT00 (Fig. 2E). There was also a higher proportion of transcripts associated with *Cell Wall Synthesis & Elongation* that are expressed only in I1 and I5 (Fig. S9). Transcripts associated with *Amino Acid Metabolism* peaked between ZT12 and ZT14 in L1 (50%). In I1 and I5, they had two peaks: between ZT00 and ZT02 (37.5% and 57.1%) and between ZT08 and ZT10 (37.5% and 42.9%) (Fig. 2B). Transcripts associated with *Transporters* peaked at ZT02 (35.7%) and ZT12 (15.7%) in L1. In I1, most of the transcripts peaked 2 h earlier, at ZT00 (24.2%) and ZT10 (24.2%). I5 displayed a similar pattern to I1, with 53.6% peaking between ZT00 and ZT02 and 46.4% between ZT08 and ZT10 (Fig. 2B). This tendency for L1 to have later phases than I1 and I5 can be seen in the putative ortholog *SWEET2,* which peaked at ZT02 in L1 and at ZT18-20 in I1 and I5 (Fig. S9L).

### Circadian clock transcripts have similar dynamics in different sugarcane organs

The differences in the rhythmic transcripts of the three organs could be explained by the presence of organ-specific circadian clocks that could generate different patterns of rhythmic transcription. For this reason, we looked at rhythms in the *Input Pathways, Central Oscillator,* and *Output* Pathways of the circadian clock. Most of the known *Input Pathways* to the circadian clock are associated with *Light Signaling^6^. Light Signaling* is underrepresented among the transcripts expressed in I5 and the rhythmic transcripts in I1 (Fig. S8). Among the red light receptor genes, *PHYTOCHROME A.1 (PHYA.1)* was rhythmic in L1, with a peak at ZT23, while *PHYB* was not rhythmic in any organ (Fig. S10A, B). In I1 and I5, both *PHYs* had two peaks: one near dawn (ZT00-02) and another at night (ZT18-20). Among the blue light receptors, *CRYPTOCHROME1.1 (CRY1.1)* was rhythmic in L1, peaking at ZT03. *CRY2.1* was rhythmic in I5, peaking at ZT19. *ZEITLUPE (ZTL.1)* was rhythmic in L1 and I5, peaking at ZT01 and ZT21, respectively (Fig. S10D-F).

The transcripts associated with the *Central Oscillator* displayed rhythms with similar dynamics (Fig. 3). *LHY* peaked early in the morning, between ZT02 and ZT04, with overlapping dynamics in all three organs (Fig. 3a). Similarly, *TOC1* peaked around dusk, between ZT10 and ZT12, in all three organs (Fig. 3D). The normalizations used to analyze the oligoarray data do not allow the comparison of expression levels, so we used RT-qPCR to show that *LHY* varied during the day by 750× in L1 and 150× in I1 and I5 (Fig. S11S). In contrast, *TOC1* differed 30× in L1 and 18× in I1 and I5 (Fig. S11B). The other PRR genes, *PRR59, PRR73,* and *PRR95* (referred to as *ScPRR3, ScPRR7,* and *ScPRR59,* respectively, in Hotta *et al.,* 2011), peaked between ZT06 and ZT10 (Fig. 3B, C, and E). *GI* peaked between ZT08 and ZT10 in all three organs (Fig. 3F). Finally, *ELF3* was rhythmic only in L1, with a peak at ZT14. In the internodes, *ELF3* had a similar pattern, but it was not regarded as rhythmic due to high noise (Fig. S10C).

**Figure 3 –.**
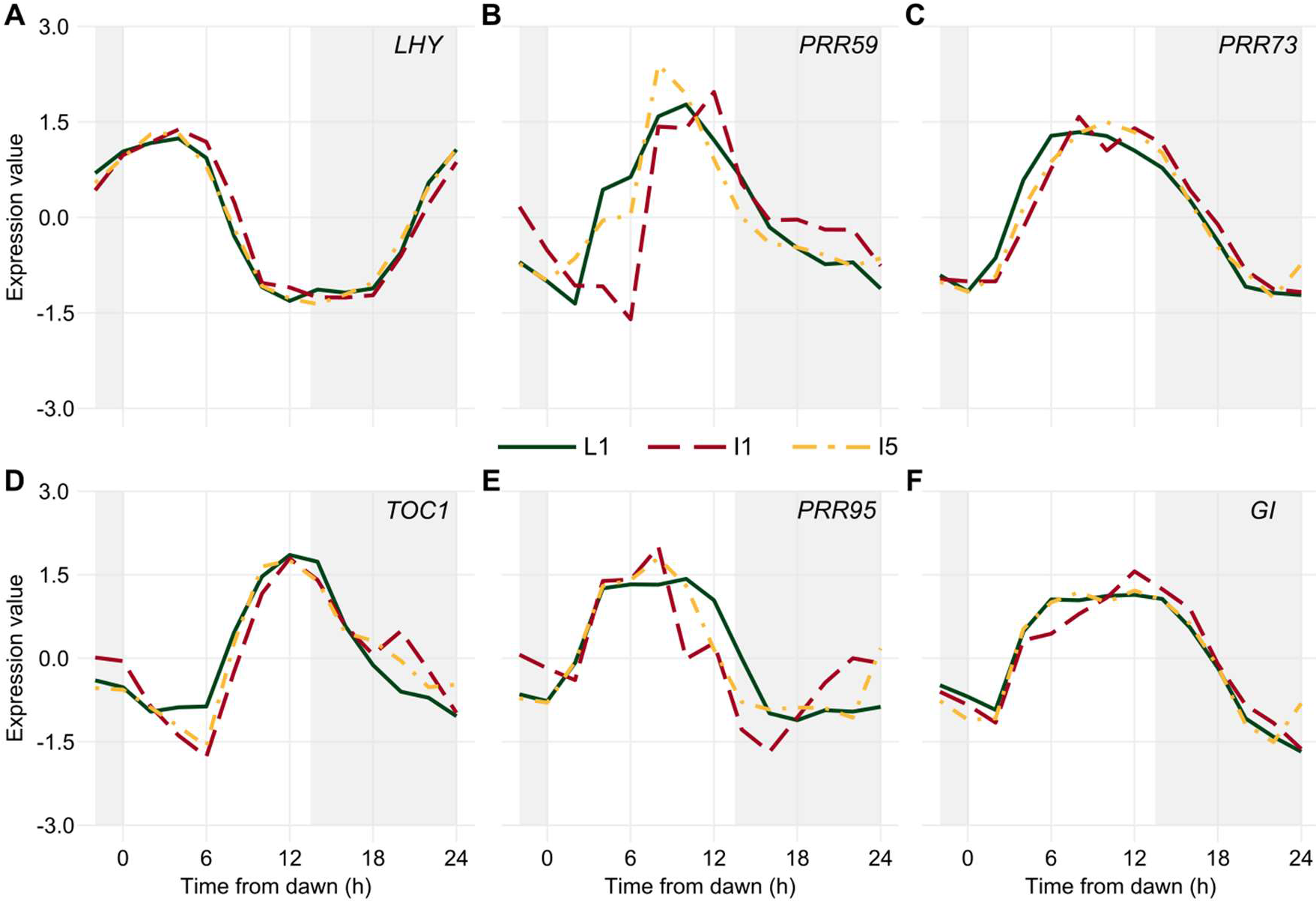
Diel rhythms of *Central Oscillator* transcripts in sugarcane organs. *LHY* (A), *PRR59* (B), *PRR73* (C), *TOC1* (D), *PRR95* (E), and *GI* (F) rhythms were measured in leaf +1 (L1, green continuous line), internodes 1 and 2 (I1, red dashed line), and internode 5 (I5, yellow dash-dotted line) of field-grown sugarcane using oligoarrays. Time series were normalized using Z-score. The light-gray boxes represent the night periods.

Among the possible pathways that can be recruited by the circadian clock that are considered part of the *Output Pathways* are those associated with *Chromatin Remodeling, Transcription Factors,* and *Protein Synthesis* (Fig. 4). Transcripts associated with *Chromatin Remodeling* peaked at ZT00-02 and ZT10-12 in L1 (32.5% and 36.5%, respectively). In I1 and I5, they peaked at ZT00 (40.7% and 40.8%, respectively) and ZT08-10 (33.99% and 51.2%, respectively) (Fig. 4A). Transcripts associated with *Transcription Factors* tended to peak near dawn, at ZT00-02, in all three organs (57.5% in L1, 46.4% in i1, and 50.3% in I5). A higher proportion (22.6%) of transcripts associated with *Transcription Factors* were rhythmic when compared to all rhythmic (16.6%) transcripts, χ^2^(6, *n* = 341) = 15.1, *P* = 0.02 (chi-square test, Fig. 4E). These transcripts also peaked similarly in all organs: 79.3% peaked in the same interval in L1 as in I1, 72.2% peaked in the same interval in L1 and I5, and 93.1% in I1 and I5.

**Figure 4 –.**
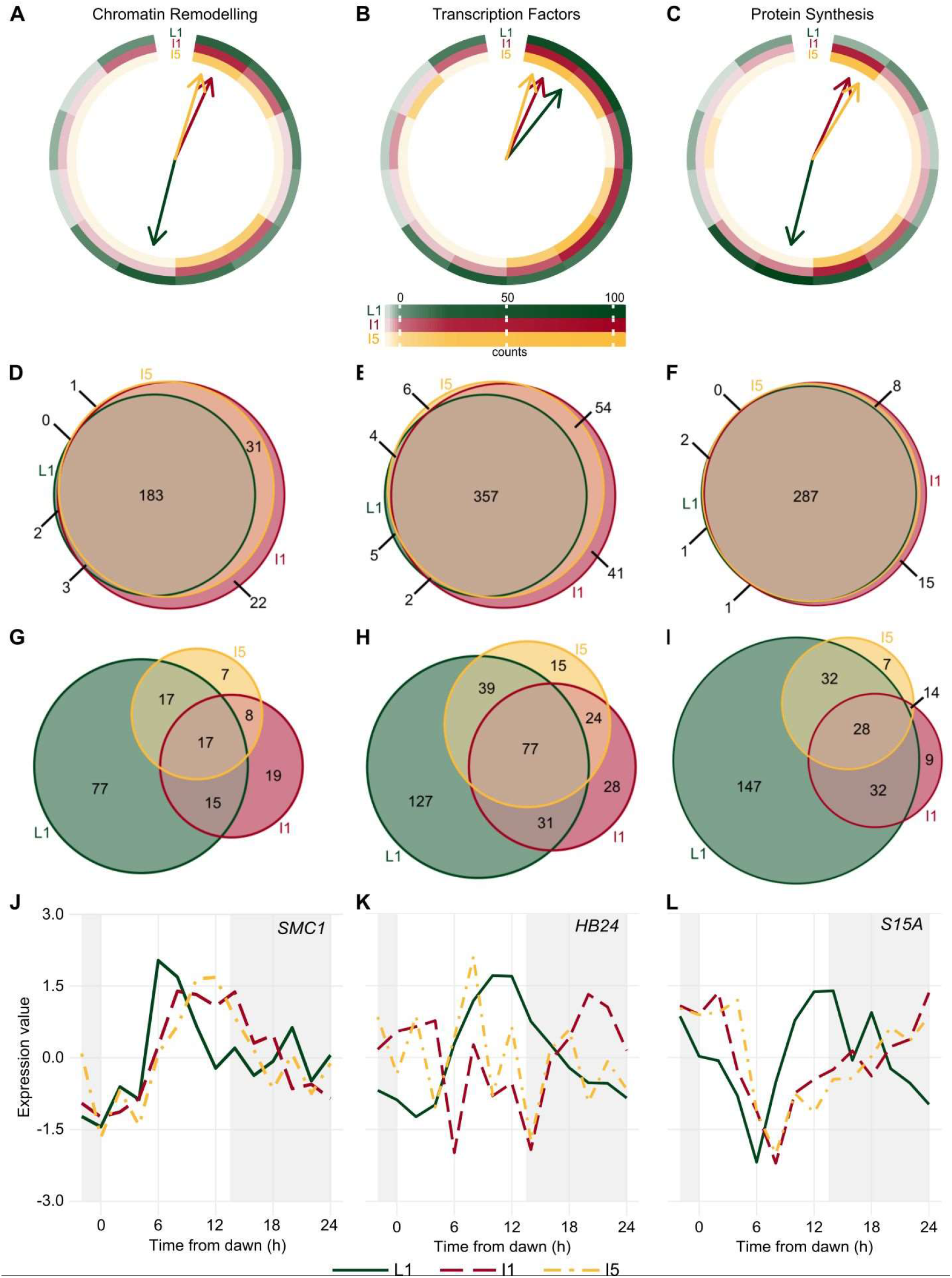
Transcripts associated with *Genetic Information Processing* have different rhythms in sugarcane organs. **(A-C)** Circular heatmap of the distribution of the peak time of rhythmic transcripts related to *Chromatin Remodeling* **(A)**, *Transcription Factors* **(B)**, and *Protein Synthesis* **(C)** in leaf +1 (L1, green), internodes 1 and 2 (I1, red), and internode 5 (I5, yellow). The colored arrows show the time at which the most transcripts are found in each organ. **(D-I)** Euler diagrams of all expressed transcripts **(D-F)** and rhythmic transcripts **(G-I)** in L1, I1, and I5 in field-grown sugarcane in diel conditions. (J-L) *SMC1* **(J)**, *HB24* (K), and *S15A* **(L)** rhythms measured in L1 (continuous green line), I1 (red dashed line), and I5 (yellow dash- dotted line) of field-grown sugarcane using oligoarrays. Time series were normalized using *Z*-score. The light-gray boxes represent the night periods.

Transcripts associated with *Protein Synthesis* tended to peak at dusk in L1 (ZT12, 49.0%), at dawn and afternoon in I1 (ZT00, 36.1%; ZT10, 32.5%), and at dawn in I5 (ZT00, 61.7%) (Fig. 4C). A high proportion of transcripts associated with *Protein Synthesis* were expressed in all three organs (91.4%) (Fig. 4F). In contrast, more than half of the transcripts (54.6%) were rhythmic only in L1, whereas a lower frequency (41.5%) of total rhythmic transcripts were seen only in L1, χ^2^(6, *n* = 269) = 34.8, *P* < 0.001 (Fig. 4I).

Other transcripts showed a wide variety of oscillations amongst the three organs (Fig. 4J-L, and S11). The putative *STRUCTURAL MAINTENANCE OF CHROMOSOMES1 (SMC1),* associated with *Chromatin Remodeling,* peaked at ZT06 in L1 and ZT11 in I1 and I5 (Fig. 4J). Two putative *JUMONJI-C (JMJC) DOMAIN-CONTAINING PROTEIN5 (JMJD5)* genes, encoding proteins that can act as histone demethylases, were found in sugarcane. *JMJD5.1* is expressed only in I1 and I5 and has a phase at ZT10 (Fig. S12A, D); *JMJD5.2* is expressed in all organs with similar rhythmic patterns (Fig. S12A). The transcription factor gene *HOMEOBOX PROTEIN24 (HB24)* is rhythmic only in L1, with a peak at ZT10 (Fig. 4K). Another rhythmic gene, *40S RIBOSOMAL PROTEIN S15 (S15A),* associated with *Protein Synthesis,* has a peak at ZT14 in L1 and at ZT00 in I1 and I5 (Fig. 4L).

## Discussion

Organ-specific rhythms of transcription can be found in highly productive and intensively selected commercial sugarcane. The specialization of the rhythmic transcriptome may help the plant cells to adapt to local environmental rhythms, as well as to generate rhythms that are compatible with their specialized needs. Specialized rhythms may also be essential to rhythmic processes that require organ-to-organ coordination, such as sucrose transport from the leaves to the internodes^28^.

### Rhythms in field conditions are different from those in controlled conditions

Sugarcane leaves in field conditions had twice as many transcripts identified as rhythmic than plants assayed under circadian conditions. This difference is expected because some rhythms are driven by environmental oscillations, such as light and temperature. Also, some circadian-clock-driven rhythms may undergo amplitude increases due to a general increase in the amplitude of the *Central Oscillator.* In L1, the transcriptional rhythms of *LHY* vary by up to 60× in a day in circadian conditions and 750× in field conditions, while those of *TOC1* vary up to 5× in a day in circadian conditions and 40× in field conditions (Hotta et al., 2013, and Fig. S11).

In circadian conditions, most transcripts peaked at subjective dusk (ZT12, 29.0%), which resulted in 60.5% of the transcripts peaking during subjective night. By contrast, in field conditions, most transcripts peaked near dawn (ZT00-02, 35.2%) in L1, which resulted in 80.3% of the transcripts peaking during the day. This reinforces the role of the light/dark transition as the driving force of rhythms in leaves in field conditions. A high proportion (64.1%) of the transcripts that peaked during the subjective night in circadian conditions showed phase changes that made their peak happen during the day in field conditions. This might suggest the existence of dampening mechanisms that actively decrease nocturnal peaks. A similar mechanism keeps cytoplasmic calcium concentration lower during the night under diel conditions (day/night) than during the subjective night under circadian conditions^29^. Most of the transcripts associated with the *Central Oscillator* maintained their core phases, except *LHY,* which had a later peak (ZT01 in circadian conditions; ZT04 in field conditions). As a comparison, *LHY* is induced by light in Arabidopsis and is mostly insensitive to temperature in rice^22,30^. In sugarcane, alternative splicing of LHY correlates with environmental temperature^31^. The differences between the rhythmic transcriptomes in circadian conditions and field conditions highlight the importance of experiments done under field conditions to understanding how the circadian clock can affect the plant transcriptome *in natura*. For example, simulations of natural conditions in growth chambers showed that the flowering signal *FLOWERING LOCUS T (FT)* has a different phase under such conditions than it does under controlled conditions in Arabidopsis^19^. This discovery will require adjustments to the current flowering models to reflect events in natural conditions.

In recent years, the productivity gains of sugarcane crop through classical breeding has been decreasing^32,33^. A possible strategy to increase productivity gains is the use of molecular markers^33–35^. However, the association between genotype and phenotype remains a challenge, despite many attempts^36,37^. Several studies have identified drought-induced genes in order to identify targets for molecular breeding^25,38–40^. However, as most of these studies only harvest at one timepoint, it is possible that important rhythmic drought-induced genes are missed^41^. In addition, delays in the harvesting of plant material, or changes in phase or period of rhythmic genes, can lead to the genes to be incorrectly considered differentially expressed^42^. Thus, the identification of rhythmic genes in the field can both increase the identification of genes of interest and help to reduce the number of false positive, aiding the identification of targets for molecular breeding.

### Rhythmic transcripts are organ-specific

The transcripts in L1, I1, and I5 have very different rhythmic patterns, even though most of the expressed transcripts were found in all three organs. Rhythms in I1 and I5 were more like each other than to those in L1, and only 7.4% of the transcripts expressed in all three organs showed the same rhythms. Thus, we conclude that these three organs have vastly different and specialized circadian clocks. These specialized circadian clocks could be the result of multiple organ sensitivities to environmental cues, of organ-specific *Core Oscillators,* and of organ-specific interactions of *Output Pathways* with environmental signals^43,44^.

In Arabidopsis, different sensitivities to environmental cues are found in the vascular phloem companion cells, which are more sensitive to photoperiodism, and the epidermis cells, which are more sensitive to temperature^43^. In sugarcane, most L1 transcripts peak at ZT00-02 and ZT12, following dawn and dusk, while most I1 and I5 transcripts peak at ZT00 and ZT08, following dawn and the daily light and temperature maxima. Thus, the circadian clocks of these organs respond differently to environmental cues such as photoperiod, light/dark transition, or temperature, as in Arabidopsis. In rice, a significant proportion of rhythmic transcripts are regulated either by the circadian clock or by temperature oscillations^22^. In sugarcane, rhythmic L1 transcripts that were also rhythmic in circadian conditions had peaks that follow *LHY* or *TOC1* expression. On the other hand, rhythmic L1 transcripts that were not rhythmic in circadian conditions peaked at dawn and dusk. In internodes, transcripts peaked at dawn and at the light and temperature maxima. Such organ-specific sensitivity to environmental cues was previously described in the vasculature and leaf epidermis^43^.

The *Central Oscillators* of mesophyll and vasculature in Arabidopsis have similar components but with different amplitudes. *AtELF4* rhythms have an amplitude 10× higher in the vasculature, *AtPRR7* and *AtPRR9* amplitudes are 2× higher in the mesophyll, and *AtTOC1* amplitude is analogous in both tissues^45^. In sugarcane, *LHY* amplitude is 6× higher and *TOC1* amplitude is 2× higher in L1 compared to I1 and I5. As leaves are exposed to direct sunlight, whereas internodes are protected by layers of leaf sheaths, it is probable that sunlight is responsible for these amplitude differences. The dynamics of *LHY, TOC1,* and *GI* during the day were very similar in the three organs. Indeed, they were considered to be coexpressed when analyzed together (data not shown). As the three organs have different levels of exposure to the environment, the existence of a common environmental signal is unlikely. Alternatively, the oscillators of the three organs could be coupled. There is evidence in Arabidopsis of root oscillators being regulated by the oscillators of the aerial parts of the plants, either the leaves or the shoot apical meristem (SAM)^46,47^. As the leaves are a source signal to both internodes, it is possible that synchronizing signals are transported with sucrose and other sugars. In Arabidopsis, sugars can also act as an entrainment signal^7,48^.

Even though there is much evidence for tissue-specific circadian clocks in Arabidopsis^29,45,49–51^, less is known about their effect on the rhythmic regulation of transcripts. In contrast, tissue-specific rhythms have been widely studied in mammals^52–55^. Sampling of 12 different mouse organs over time showed that 43% (~8,500) of all transcripts had circadian rhythms in at least one organ, but only 10 transcripts were rhythmic in all organs^54^. As in sugarcane leaves, the rhythmic transcripts in mammalian organs tended to peak at dawn and dusk. In general, the only transcripts that had similar phases across all organs were the ones associated with the mammalian *Core Oscillator*^54^

At least two regulatory pathways are required to generate tissue-specific sets of rhythmic transcripts: one that confers organ specificity and one that confers rhythmicity. These pathways can be organized in different nonexclusive ways: they could act on a gene independently, the tissue specificity pathways could regulate the rhythmicity pathways, or the rhythmicity pathways could regulate the organ specificity pathways (Fig. S13). The rhythmicity pathways can be dependent on the circadian clock, on environmental rhythms, or both. The tissue specificity pathways can include transcription factors, protein-protein interactions, alternative promoter usage, and chromatin interactions^56^.

In our datasets, transcripts that were expressed only in one organ or only in the internodes were less likely to be considered rhythmic (Fig. 2B). Thus, it is possible that rhythmic pathways regulate only a small proportion of organ-specific pathways. Transcripts associated with *Transcription Factors* were more likely to be rhythmic in all three organs, and these transcripts had a higher probability of having the same phase. However, just a few tissue-specific rhythms in transcription factors can have a sizeable cascading effect^57^. Tissue-specific transcription factors, even if nonrhythmic, could also change the phase of rhythmic transcripts through protein-protein interactions or by changing the promoter usage^56,58^. Finally, chromatin remodeling could be a significant regulatory pathway in the generation of the tissue-specific rhythmic transcriptome. In Arabidopsis, chromatin remodeling can regulate the *Central Oscillator,* but little is known about how the plant circadian clock can use chromatin remodeling to generate rhythms^59–62^. In sugarcane, rhythms in transcripts associated with *Chromatin Remodeling* were underrepresented among the rhythmic transcripts. However, chromatin remodeling tends to be regulated post-transcriptionally through histone modifications. Transcription can also be regulated at the chromatin level through topologically associating domains (TADs). TADs are domains of DNA that self-interact, generating regulatory compartments within the chromosomes^63^. An enhancer only interacts with a gene if they share the same TAD. In consequence, it is possible to change the enhancers that interact with a gene by changing the boundaries of a TAD, which are maintained by cohesins and CCCTC-binding factors (CTCF) in mammals^63^. TADs can be regulated to generate tissue-specific transcription and even rhythms^64–66^. In plants, TADs are maintained by cohesins, but there are still no known CTCF counterparts^67,68^. In sugarcane, the cohesin subunit *SMC1* has different phases in leaves and internodes (Figure 4J).

### The role of organ-specific rhythms in sugarcane

Organ-specific rhythms may affect sugarcane productivity as they allow the different tissues to be more efficient according to their function and local environmental rhythms^56^. In mammals, rhythms in fibroblasts allow wound healing to occur faster during the active phase than the rest phase^69^. Rhythms in the liver lead to larger cell sizes and protein levels during the active phase and after feeding, making detoxification more efficient during the active and post-feeding periods^70^.

Nutrient and photoassimilate transportation inside the plant is essential for rhythmic processes and may also be part of organ-to-organ coordination and C partitioning^28,71–73^. In sugarcane, expressed transcripts associated with *Transporters* were overrepresented in L1 and I5, while rhythmic transcripts associated with *Transporters* were overrepresented in L1 and I1. Furthermore, transcripts associated with *Transporters* tended to peak 2 h later in L1 than in the internodes, which may indicate that the latter is the driving force of this process. The phloem and xylem are also important organs for the integration of multiple rhythmic information generated by specialized circadian clocks, such as flowering^74^.

Sugarcane have rhythms of sucrose and starch in the leaves but not in the internodes^75^. In this crop, sucrose is synthesized in the leaves and is degraded in the apoplast or cytosol of internodes to be re-synthetized in their vacuoles^28^. Organ- specific regulation of transcripts may regulate sucrose storage in sugarcane. Differences in the rhythms of transcripts associated with *Carbohydrate metabolism* may be a way to regulate C partitioning to different organs. In our experiments, transcripts associated with *Carbohydrate metabolism* peaked later in internodes than in the leaves (Figure 2E). *CWI* had a peak at ZT04 in L1, and at ZT08 in 11 and I5 (Figure S9Q). In sugarcane, higher activities of cell wall invertases are associated with higher sucrose content, possibly by enhancing sucrose unloading in the internodes ^28,76–78^. *SPSII,* one of the enzymes that synthesize sucrose, was only rhythmic in L1, with a morning peak (Figure S9M). Interestingly, two rhythmic *SuSy* had the same dynamics in all organs (Figure S9I) but the other four were only expressed in the internodes. Sucrose synthases can function either degrading or synthesizing sucrose. In sugarcane, *SuSys* mainly work in the degradative direction, and their pattern of expression are associated with the regulation of sucrose uptake in the internodes ^28,79,80^.

## Conclusions

The vast differences found in the rhythmic transcriptomes of different plant organs provide important clues to understanding the way that tissue-specific circadian clocks are generated and their impact on plant physiology. However, little is still known about the molecular mechanisms that control this specialization. The combination of organ- or tissue-specific studies with the observation of rhythms in the field, where conditions are fluctuating and variable as is normal in natural environments, is essential to understanding the nuances of how the plant circadian clock increases the fitness of plants and, in turn, crop productivity.

## Materials and Methods

### Plant growth and harvesting

Commercial sugarcane (*Saccharum* hybrid SP80-3280) was planted in a field in Araras, Brazil (22°18’41.0”S, 47°23′5.0″W, at an altitude of 611 m), in April 2012 (autumn) (Fig. S1). The soil on the site was a Typic Eutroferric Red Latosol. Plants were harvested 9 months later, in January 2013 (summer), after an unusually dry winter and spring. The time course experiment started 2 h before dawn and continued every 2 h until the next dawn, generating time series with 14 time points in total. Dawn was at 5:45, and dusk was at 19:00 (13.25 h light/10.75 h dark) (Fig. S1). At each time point, leaf +1 (the first leaf from the top with clearly visible dewlap), internodes 1 and 2, and internode 5 of nine individuals were harvested (Fig. S2), frozen in liquid N2, and stored in three pools of three individuals each. Two pools were used as biological replicates for oligoarrays, and one pool was used for validation using the reverse-transcription quantitative PCR (RT-qPCR).

### Oligoarray hybridizations

All frozen samples were pulverized in dry ice using a coffee grinder (Model DCG-20, Cuisinart, China). One hundred milligrams of each pulverized sample was used for extraction of total RNA using Trizol (Life Technologies), following the supplier’s instructions. The RNA was treated with 2 U DNase I (Life Technologies) for 30 min at 37°C and cleaned using the RNeasy Plant Mini kit (Qiagen). The quality and quantity of RNA were assayed using an Agilent RNA 6000 Nano Kit Bioanalyzer chip (Agilent Technologies). Sample labeling was done following the Low Input Quick Amp Labelling protocol of the Two-Color Microarray-Based Gene Expression Analysis system (Agilent Technologies). Hybridizations were done using a custom 4×44 k oligoarray (Agilent Technologies) that was previously described^15,25^. Two hybridizations were done for each time point against an equimolar pool of all samples of each organ. Each duplicate was prepared independently using dye swaps. Data were extracted using the Feature Extraction software (Agilent Technologies) (Figure S3A). Background correction was applied to each dataset. A nonlinear LOWESS normalization was also applied to the datasets to minimize variations due to experimental manipulation. Signals that were distinguishable from the local background signal were taken as an indication that the corresponding transcript was expressed. We have validated 10 transcripts (30 time series) using RT-qPCR (Figures S4 and S5). Among the time series identified as rhythmic (n = 23), 91% were also rhythmic using data from RT-qPCR (Table S2), and 77% were considered correlated using Spearman’s rank correlation coefficient. Among the time series identified as not rhythmic (n = 7), 86% were also not rhythmic using data from RT-qPCR, and 36.7% were considered correlated using Spearman’s rank correlation coefficient. The GenBank ID and Sugarcane Assembled Sequences (SAS) numbers for sugarcane genes are listed in Table S1. The complete dataset can be found at the Gene Expression Omnibus public database under the accession number GSE129543.

### Data analysis

For the purposes of further analysis, only transcripts that were found to be expressed in more than 7 of the 14 time points were considered to be expressed. All of the expressed transcripts time series were grouped in coexpressed modules using the R package weighted correlation network analysis (WGCNA) to identify rhythmic transcripts^26^ (Figure S3B). Network adjacency was calculated using a soft thresholding power of 18 for all organs. Modules that had a dissimilarity value of ≤ 0.25 were merged. Final modules were generated using a 0.175 adjacency threshold. As WGCNA groups together time series that have a positive or a negative correlation, we normalized each time series using a Z-score, separated these time series into two new modules, and generated a typical time series for each module by finding the median of all time series. Then, each representative time series was classified as rhythmic or non-rhythmic using JTK-CYCLE^27^. Modules that had an adjusted P-value of < 0.75 were considered rhythmic. Finally, we filtered out noisy time series, defined as those that had a Spearman’s rank correlation coefficient of < 0.3 when compared against the representative time series. Phase was assigned using the phase estimated by JTK-CYCLE corrected against a dendrogram with the representative time series of all modules of all organs. Modules that clustered together in the dendrogram were considered to have the same phase. The phase of a time series is defined as the time between dawn and the peak of the time course. Euler diagrams were done using the R package *eulerr.* Chi-squared (χ^2^) tests were used to compare Euler diagrams. Heatmaps were created using the R packages *circlize^81^* and *ComplexHeatmap^82^.* To evaluate if a group of transcripts were under- or overrepresented, we used a hypergeometric test *(phyper* function in R). With this test, a P-value < 0.05 suggests that the analyzed group is overrepresented in the dataset, while a P-value > 0.95 suggests that the analyzed group is underrepresented in the dataset. Code to fully reproduce our analysis is available on GitHub (https://github.com/LabHotta/sugarcane_field_rhytms) and archived on Zenodo (http://doi.org/10.5281/zenodo.2636813).

### RT-qPCR analysis

As described for the oligoarray hybridizations, 100 mg of the pulverized frozen samples for all three organs was used for total RNA extractions following the same Trizol (Life Technologies) protocol and then were treated with DNase I (Life Technologies) and cleansed using the RNeasy Plant Mini Kit (Qiagen). RNA quality and concentration of each sample were checked using an Agilent RNA 6000 Nano Kit Bioanalyzer chip (Agilent Technologies). Five micrograms of total purified RNA was enough for the reverse transcription reactions using the SuperScript III First-Strand Synthesis System for RT-PCR (Life Technologies). The RT-qPCR reactions for all samples were done using Power SYBR Green PCR Master Mix (Applied Biosystems), 10× diluted cDNA, and specific primers described by Hotta et al. (2013) (Figure S11). Reactions were placed in 96-well plates and read with the Fast 7500/7500 Real-Time PCR System (Applied Biosystems). Data analysis was performed using the Fast 7500/7500 RealTime PCR System built-in software (Applied Biosystems).

## Supporting information

All Supplemental Figures and Tables

## Acknowledgements

This work was supported by the São Paulo Research Foundation (FAPESP) (grant nos. 11/00818-8 and 15/06260-0; BIOEN Program) and by the Serrapilheira Institute (grant no. Serra-1708-16001). L.L.B.D, C. A. L., and N. O. L. were supported by FAPESP scholarships (grants 11/08897-4, 13/05301-9, and 16/06740-4, respectively). F.M.A.J was supported by CAPES scholarships (Finance Code 001).

