## Supplementary material for "Rhythms of Transcription in Field-Grown Sugarcane Are Highly Organ Specific": All Supplemental Figures and Tables

**Supporting Information (SI)**

**Supplemental Figure 1**

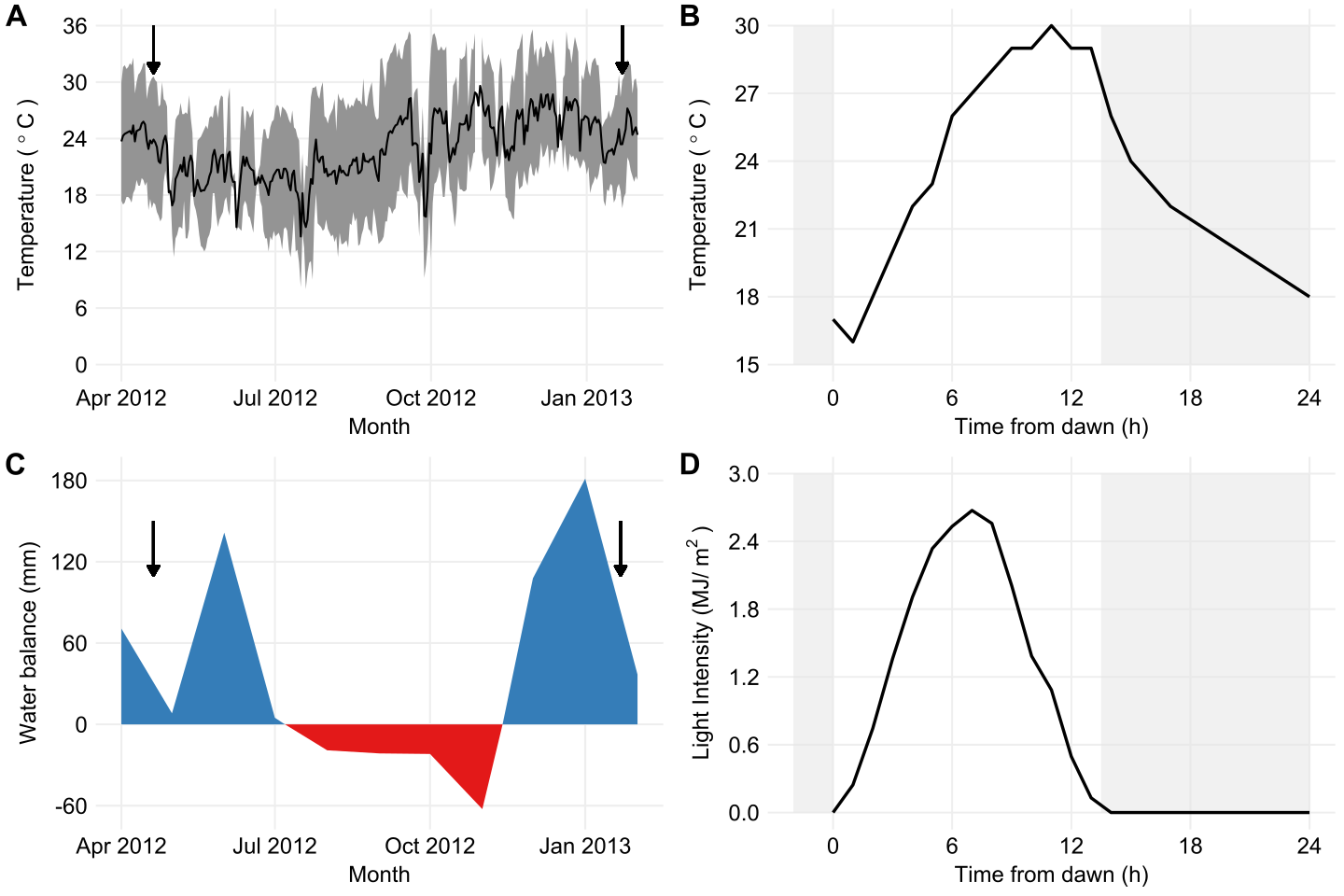

**Figure S1 – Environmental conditions in Araras (SP, Brazil).** Environmental conditions during the growth period and harvesting day. **(A)** Average temperature variation during the growth period (black line). The gray area shows the temperature variation (maximum and minimum) during each day. The arrows show the day of planting and the day of harvesting. **(B)** Temperature during the harvesting day. The light-gray boxes represent the night period. **(C)** Water balance during the growth period. **(D)** Light intensity during the harvesting day.

**Supplemental Figure 2**

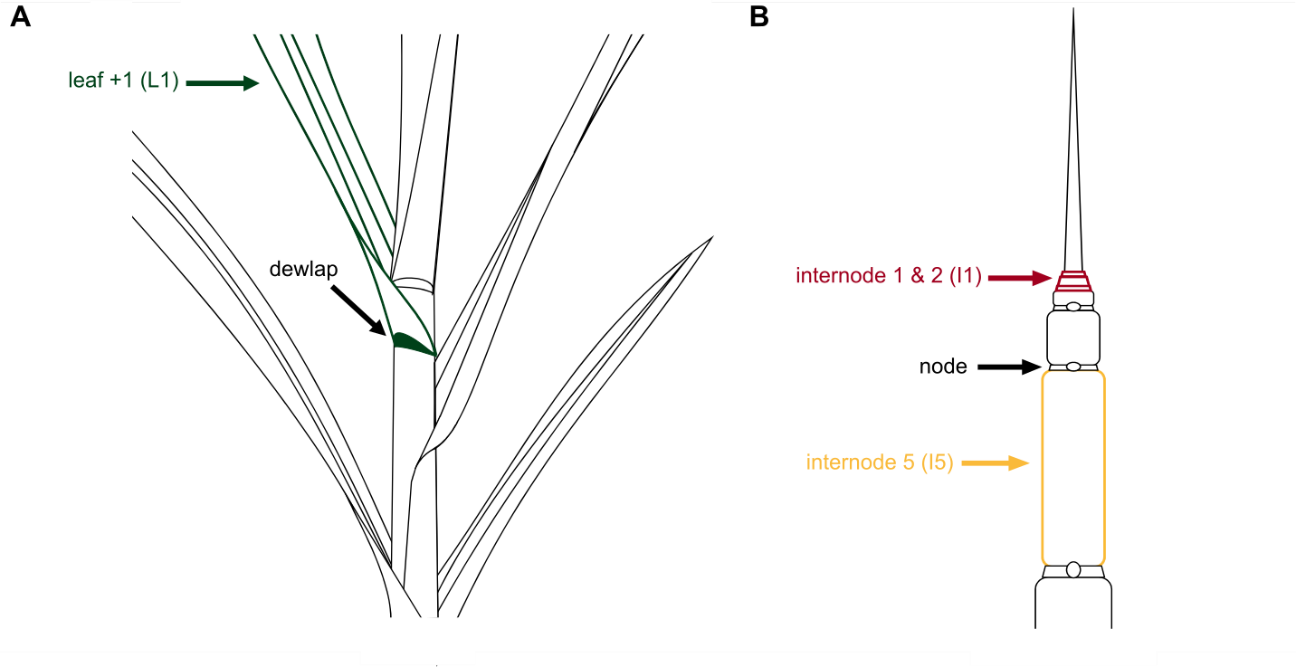

**Figure S2 – Three sugarcane organs were harvested.** **(A)** The leaf +1 (L1, green) harvested for this study is the first leaf from the top with a clearly visible dewlap. **(B)** After all of the leaves are peeled off and removed, the full stalk is exposed so that all nodes and internodes can be seen. The top two internodes (internodes 1 and 2, I1, red) and the fifth internode (internode 5, I5, yellow) were also harvested.

**Supplemental Figure 3**

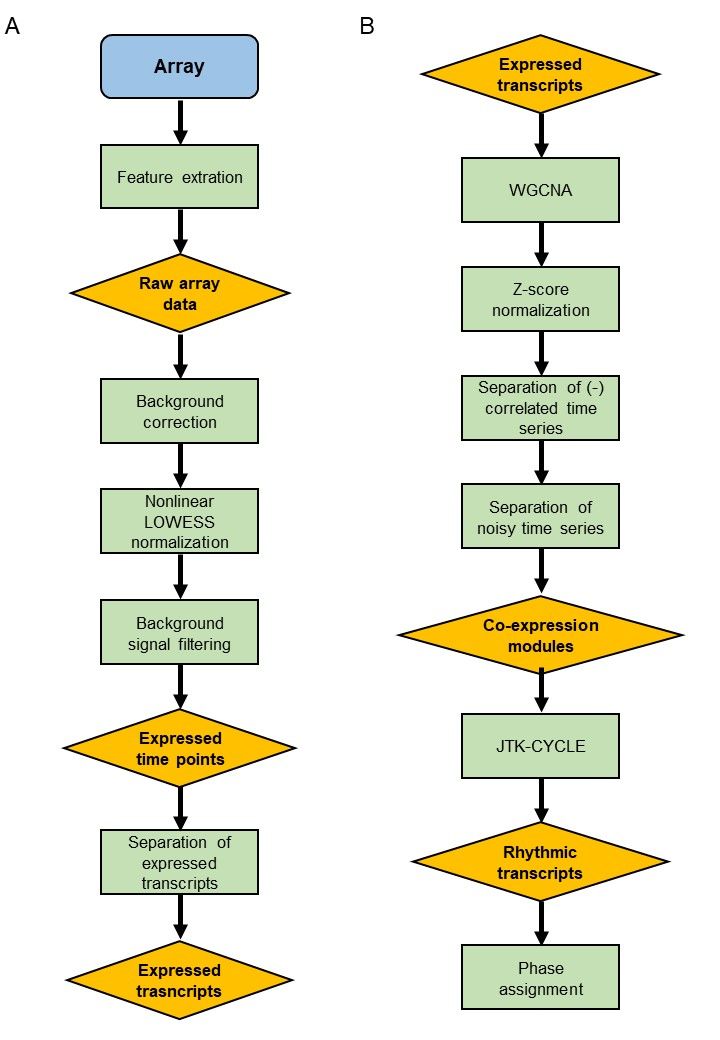

**Figure S3 – Data analysis workflow used to identify rhythmic transcripts. (A)** Workflow used to select transcripts that were expressed in > 50% of the time points of the time series. (b) Workflow used to select which expressed transcripts are also rhythmic.

**Supplemental Figure 4**

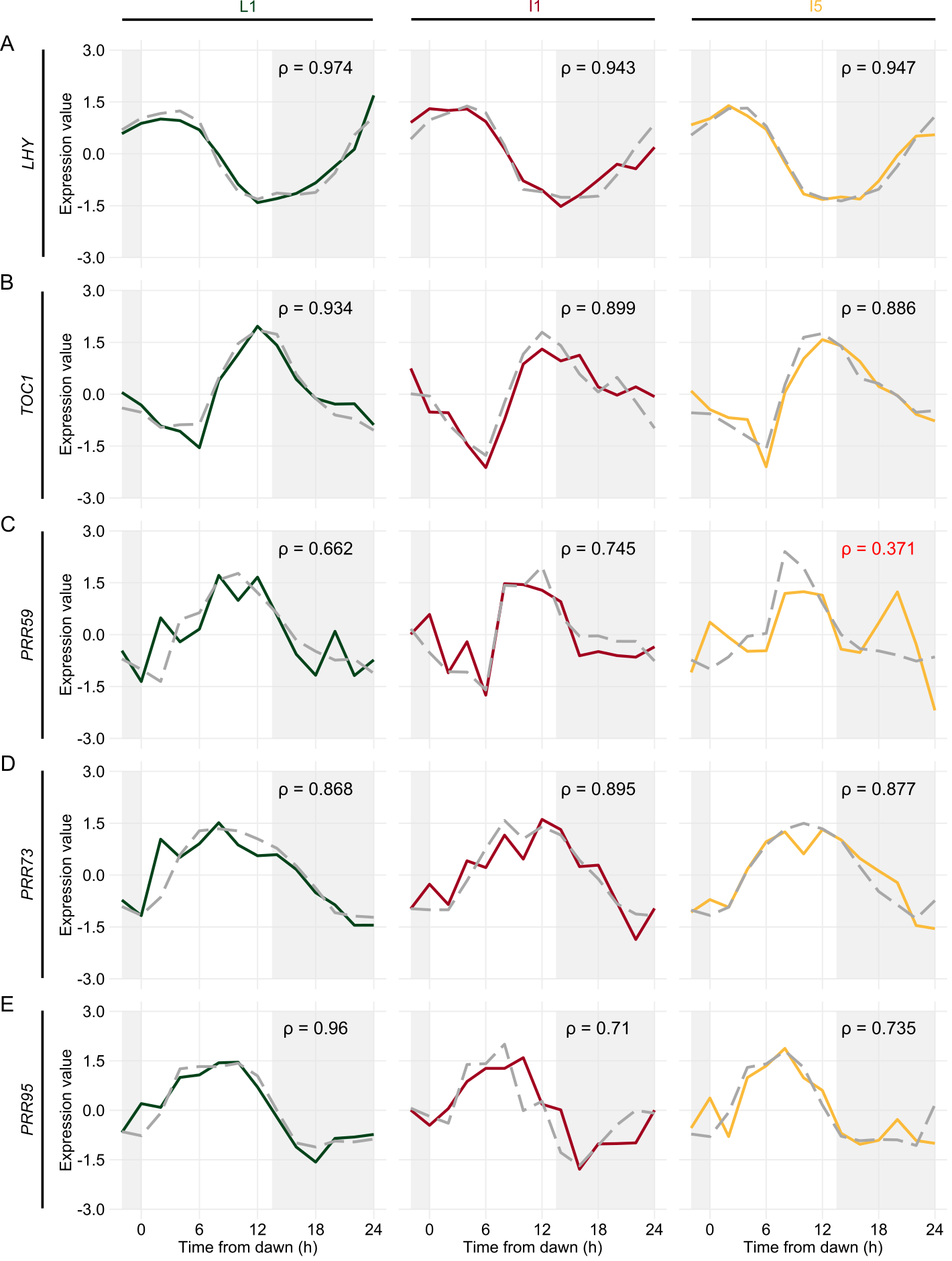

**Figure S4 – RT-qPCR validation of oligoarrays data**. Oligoarray data from leaf +1 (L1, green), internodes 1 and 2 (I1, red), and internode 5 (I5, yellow) was compared with RT-qPCR data (gray). *LHY* **(A)**, *TOC1* **(B)**, *PRR59* **(C)**, *PRR73* **(D)**, and *PRR95* **(E)**. The Spearman's rank correlation coefficient (ρ) was used to establish a positive correlation between the time series (ρ > 0.539). Time series were normalized using Z-score. The light-gray boxes represent the night period.

**Supplemental Figure 5**

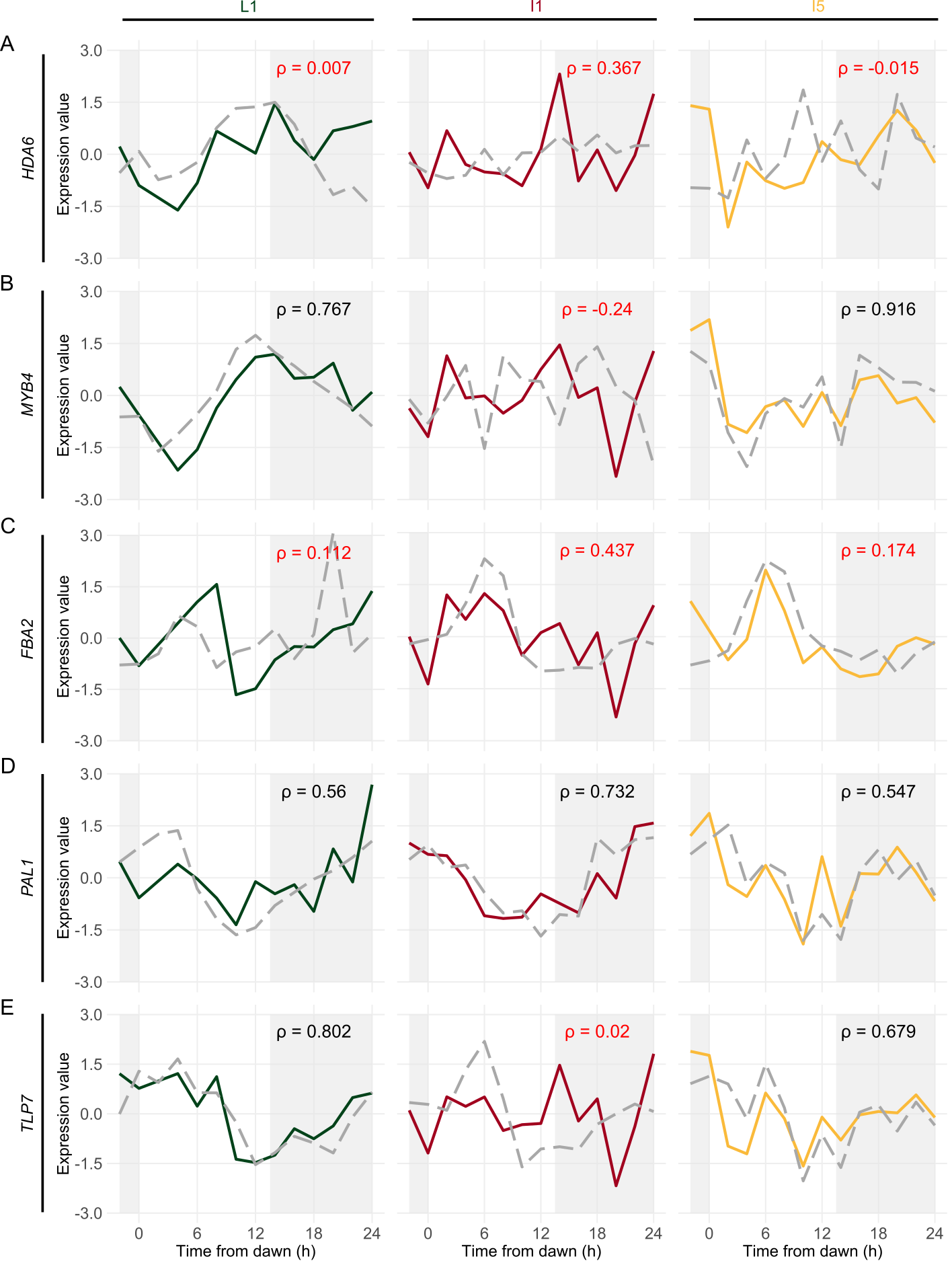

**Figure S5 – RT-qPCR validation of oligoarrays data**. Oligoarray data from leaf +1 (L1, green), internodes 1 and 2 (I1, red), and internode 5 (I5, yellow) was compared with RT-qPCR data (gray). *HDA6* **(A)**, *MYB4* **(B)**, *FBA2* **(C)**, *PAL1* **(D)**, and *TLP7* **(E)**. The Spearman's rank correlation coefficient (ρ) was used to establish a positive correlation between the time series (ρ > 0.539). Time series were normalized using Z-score. The light-gray boxes represent the night period.

**Supplemental Figure 6**

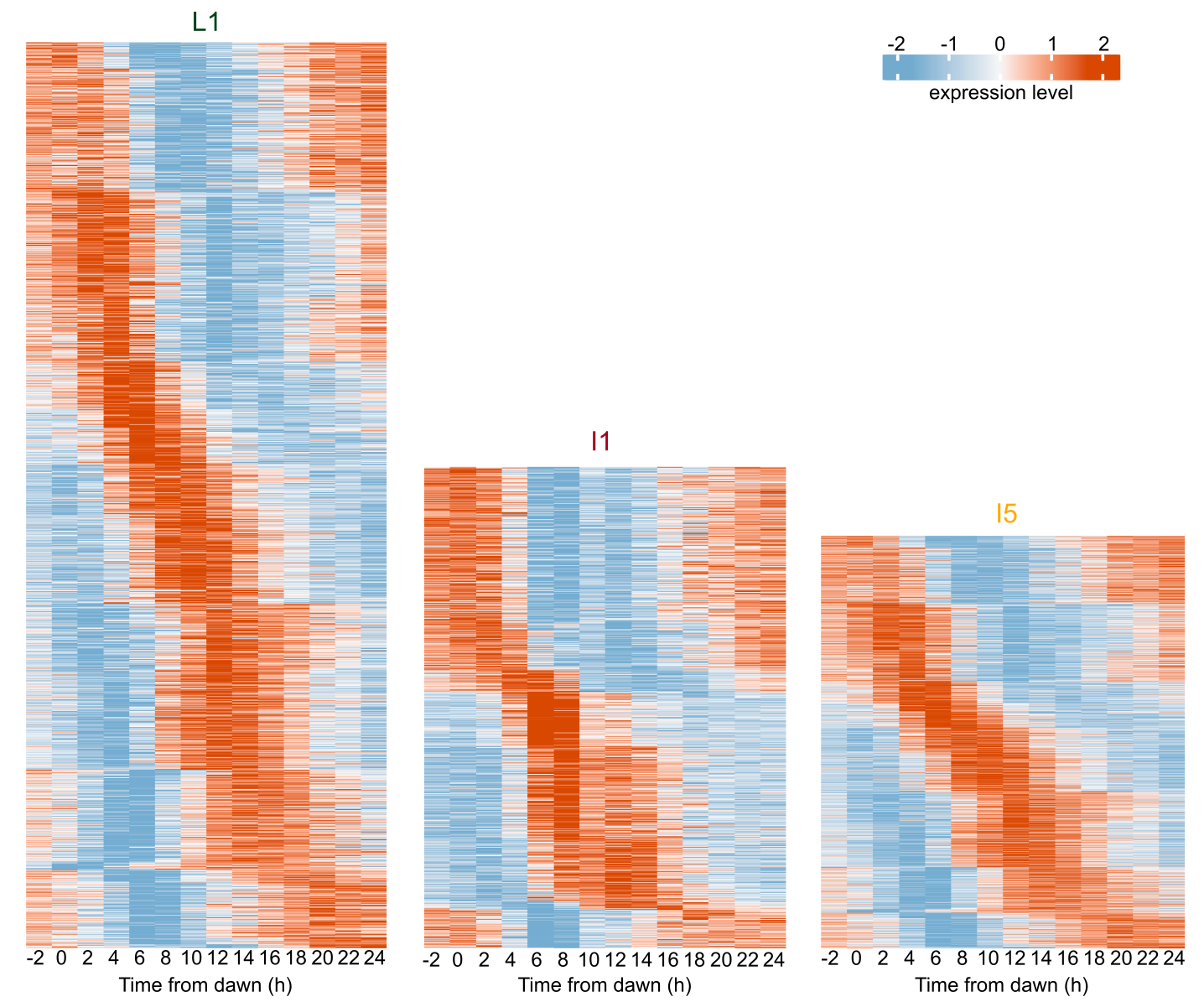

**Figure S6 – Heatmap of the expression levels of the rhythmic transcripts in sugarcane organs.** Each line represents the time course of each rhythmic transcript in leaf +1 (l1), internodes 1 and 2 (I1), and internode 5 (I5). The transcripts were separated by their phase. High transcription levels are in shades of red, and low transcription levels are in shades of blue. Transcription levels were normalized by *Z*-score.

**Supplemental Figure 7**

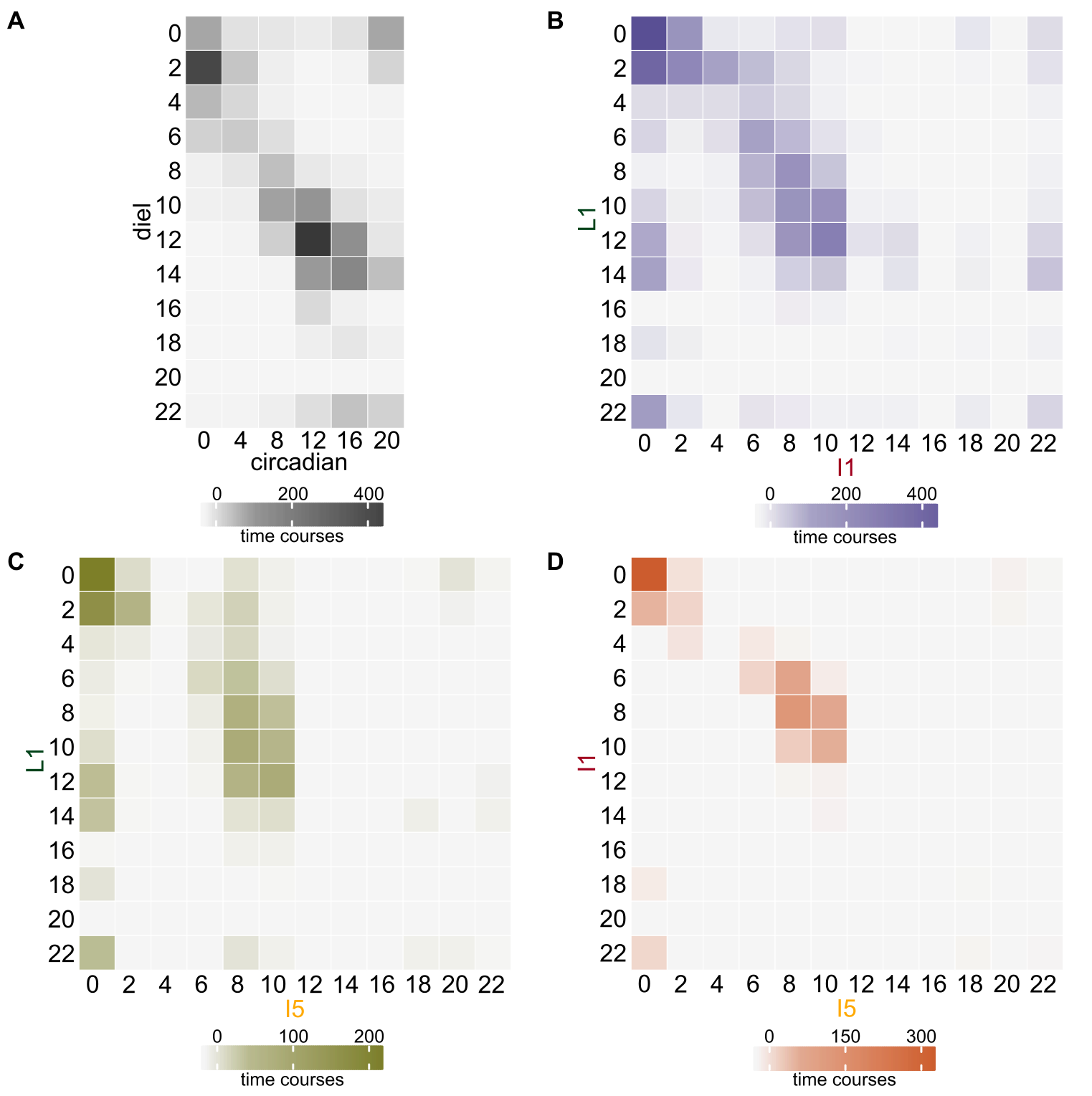

**Figure S7 – Heatmap of the phase distribution of rhythmic transcripts shared between pairs of datasets. (A)** Phase distribution of transcripts found in leaf +1 (L1) of field-grown sugarcane and L1 of sugarcane grown under circadian conditions^15^. **(B)** Phase distribution of transcripts found in L1 and internodes 1 and 2 (I1) of field-grown sugarcane. **(C)** Phase distribution of transcripts found in L1 and internode 5 (I5) of field-grown sugarcane. **(D)** Phase distribution of transcripts found in I1 and I5 of field-grown sugarcane.

**Supplemental Figure 8**

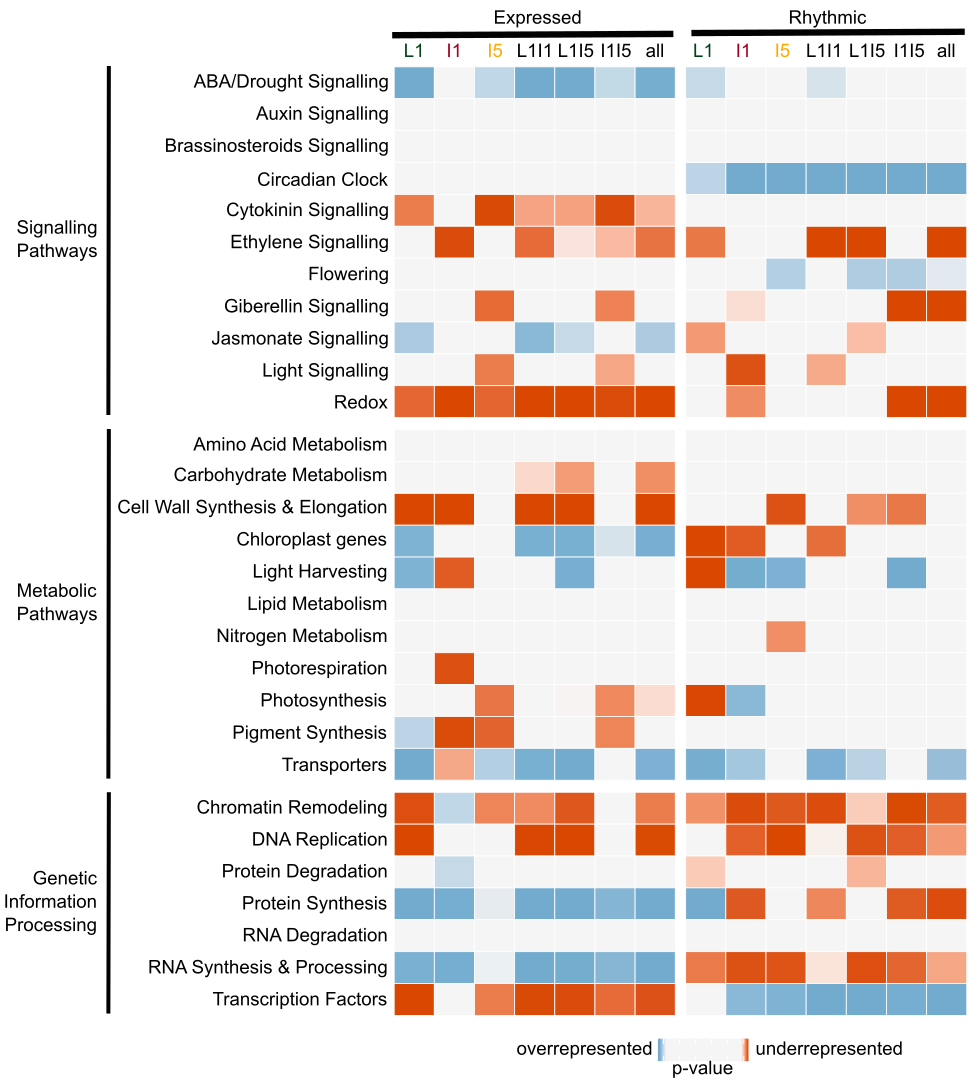

**Figure S8 –** Heatmap of functional categories that are overrepresented (shades of blue) or underrepresented (shades of red) among the expressed and rhythmic transcripts of L1, I1, I5, L1I1, L1I5, I1I5, and all three organs. The *P*-values were calculated using a hypergeometric test.

**Supplemental Figure 9**

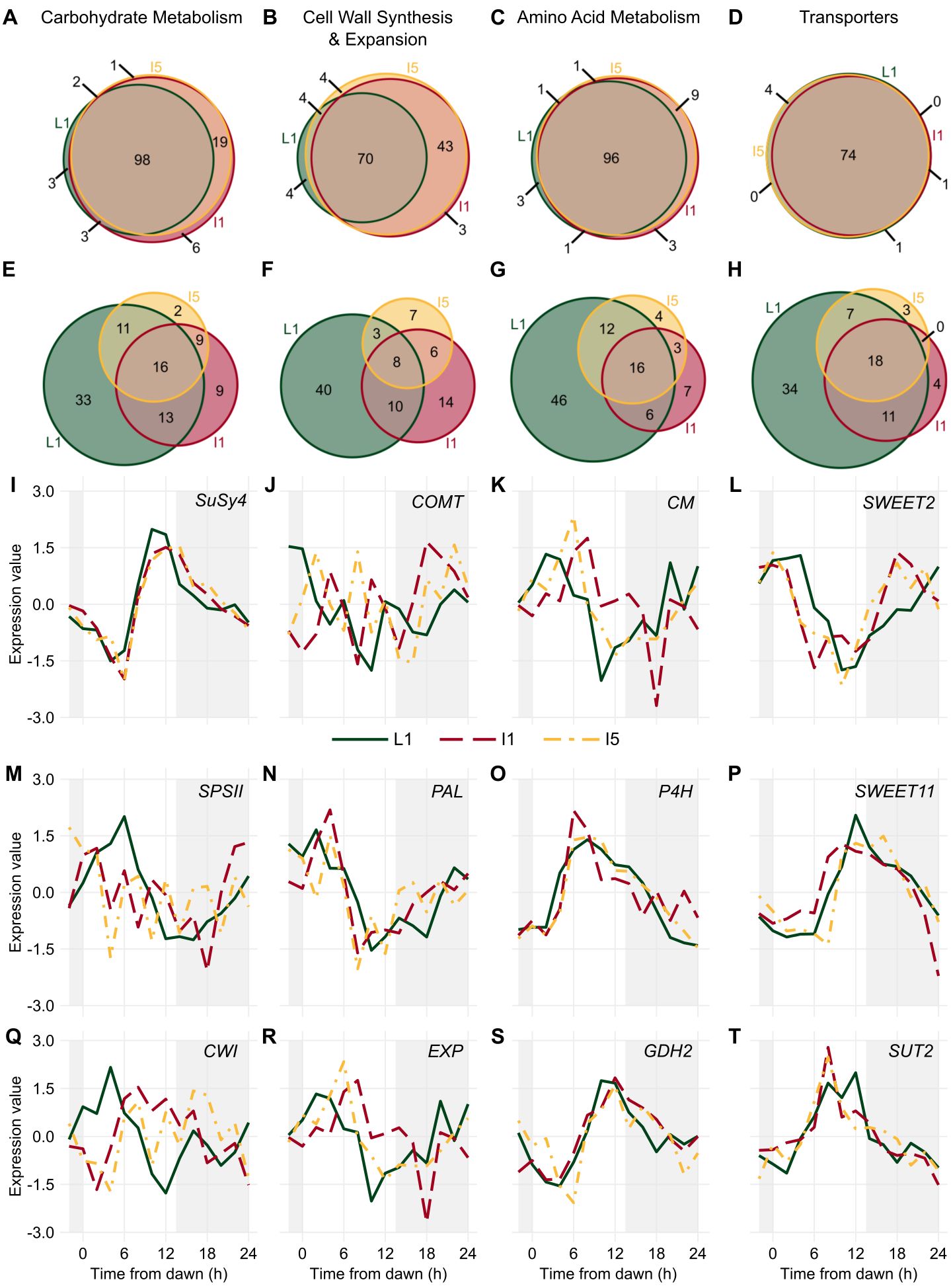

**Figure S9 – Transcripts associated with metabolic pathways have different rhythms in sugarcane organs.** **(A-H)** Euler diagram of **(A-D)** expressed and **(E-H)** rhythmic transcripts in leaf +1 (L1, green), internodes 1 and 2 (I1, red), and internode 5 (I5, yellow) in field-grown sugarcane in diel conditions. **(I-T)** Rhythms of **(I)** *SUCROSE SYNTHASE4* (*SuSy4*), **(J)** *CATECHOL-O-METHYLTRANSFERASE* (*COMT*), **(K)** *CHORISMATE MUTASE* (*CM*), **(L)** *SWEET2*, **(M)** *SUCROSE-PHOSPHATE SYNTHASE II* (*SPSII*), **(N)** *PHENYLALANINE AMMONIA LYASE* (*PAL*), **(O)** *PROLYL 4-HYDROXYLASE* (*P4H*), **(P)** *SWEET11*, **(Q)** *CELL WALL* *INVERTASE* (*CWI*), **(R)** *EXPANSIN* (*EXP*), **(S)** *GLUTAMATE DEHYDROGENASE2* (*GDH2*), and **(T)** *SUCROSE TRANSPORT PROTEIN2* (*SUT2*) measured in the L1 (continuous green line), I1 (red dashed line), and I5 (yellow dash-dotted line) of field-grown sugarcane by oligoarrays. Time series were normalized using *Z*-score. The light-gray boxes represent the night period.

**Supplemental Figure 10**

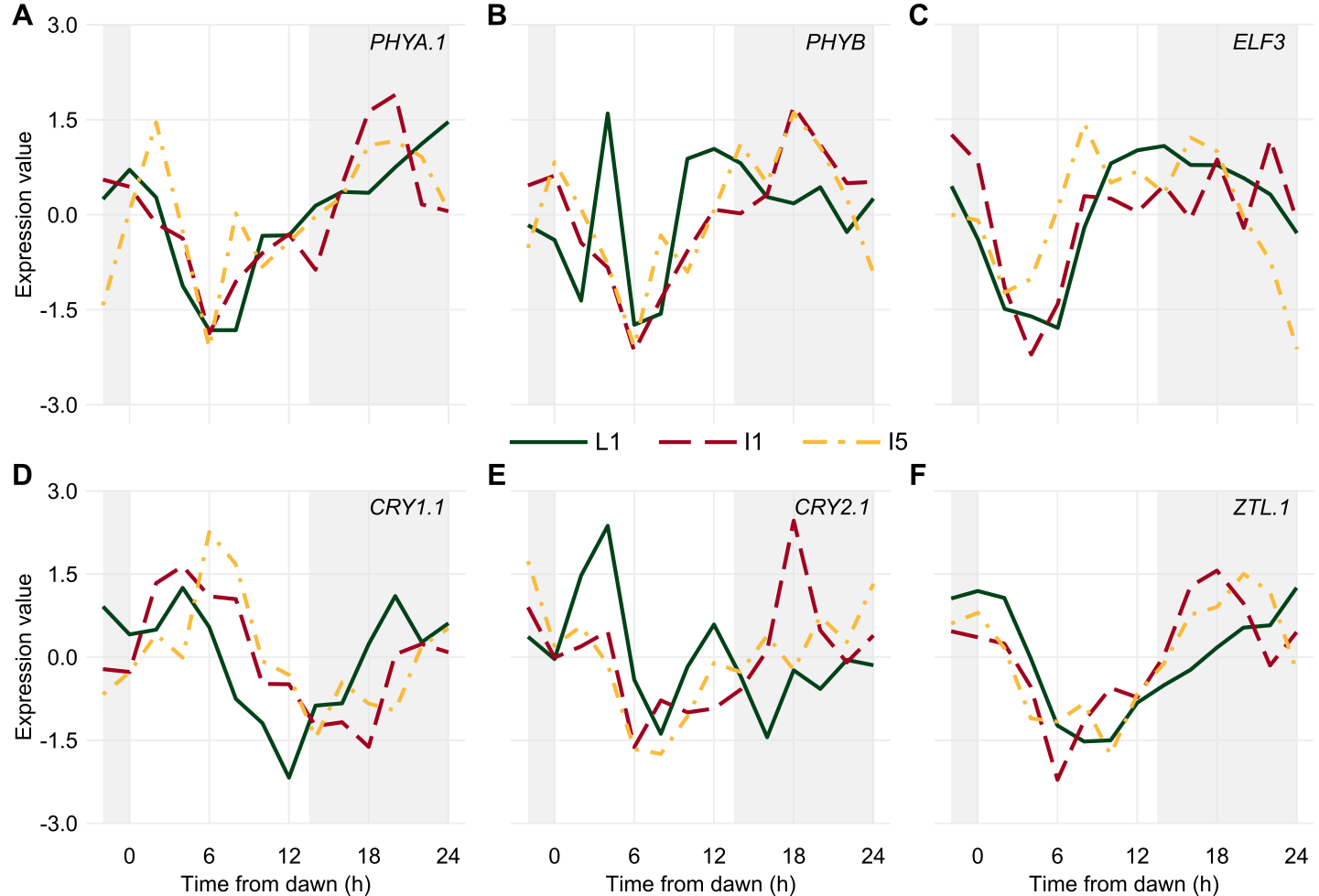

**Figure S10 – Diel rhythms of *Light Signalling* and *Central Oscillator* transcripts in sugarcane organs.** Rhythms of **(A)** *PHYTOCHROME A.1* (*PHYA.1*), **(B)** *PHYB*, **(C)** *EARLY FLOWERING3* (*ELF3*), **(D)** *CRYPTOCHROME 1.1* (*CRY1.1*), **(E)** *CRY2.1*, and **(F)** *ZEITLUPE* (*ZTL*) measured in leaf +1 (L1; continuous green line), internodes 1 and 2 (I1; red dashed line), and internode 5 (I5; yellow dash-dotted line) of field-grown sugarcane using oligoarrays. Time series were normalized using *Z*-score. The light-gray boxes represent the night period.

**Supplemental Figure 11**

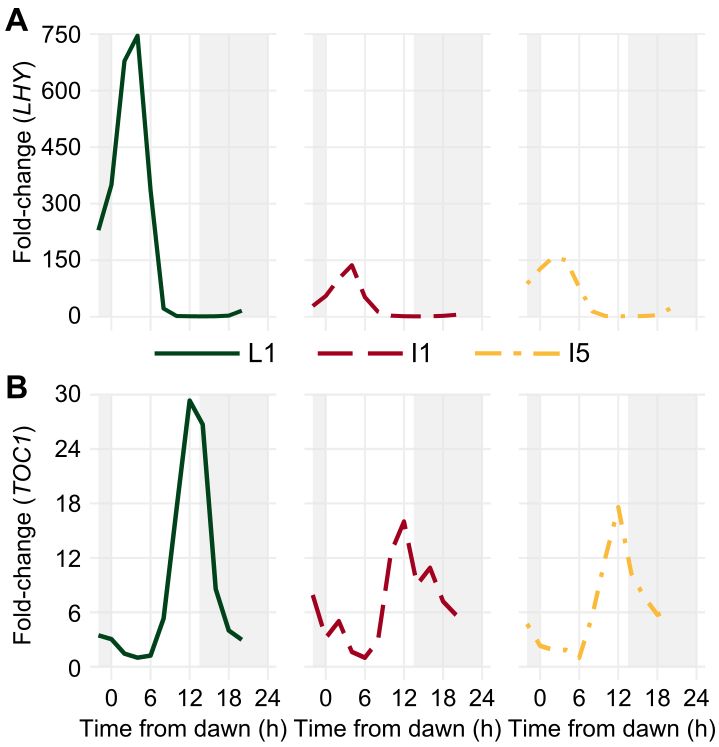

**Figure S11 – Diel rhythms of transcripts associated with the circadian clock have different amplitudes in different sugarcane organs.** Rhythms of **(A)** *LATE ELONGATED HYPOCOTYL* (*LHY*) and **(B)** *TIME OF CAB EXPRESSION1* (*TOC1*) were measured in the leaf +1 (L1, continuous green line), internodes 1 and 2 (I1, red dashed line), and internode 5 (I5, yellow dash-dotted line) of field-grown sugarcane measured using RT-qPCR. *GLYCERALDEHYDE-3-PHOSPHATE DEHYDROGENASE* (*GAPDH*) was used as a reference gene. Time series were normalized by their minimum value (minimum value = 1) using *Z*-score. The light-gray boxes represent the night period. *LHY* primers: FWD 5′-CCACCACGGCCTAAAAGAAA-3′, RVS 5′-TGGTTTTGTTGACTTGTCATTTGG-3′; *TOC1* primers: FWD 5′-TTCTGCCTGAATTTGGCAAGTG-3′, RVS 5′-GGCATCGAGCACACCAATGC-3′; *GAPDH* primers: FWD 5′-GGCATCGAGCACACCAATGC-3′, RVS 5′-TCCTCAGGGTTCCTGATGC-3′.

**Supplemental Figure 12**

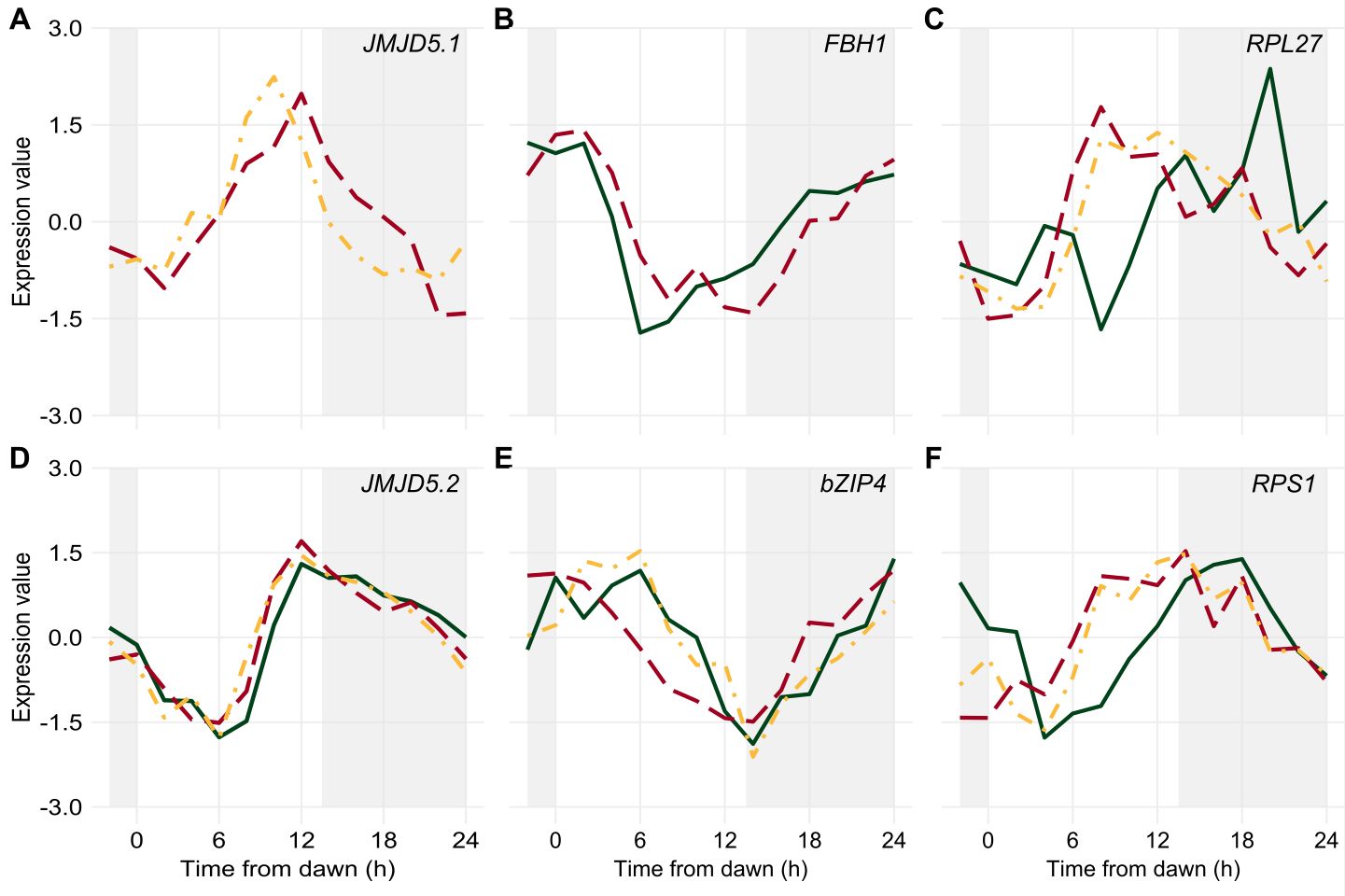

**Figure S12 – Diel rhythms of *Output Pathways* transcripts in sugarcane organs.** Rhythms of **(A)** *JUMONJI-C (JMJC) DOMAIN-CONTAINING PROTEIN 5.1* (*JMJD5.1*), **(B)** *FLOWERING BASIC HELIX-LOOP-HELIX1* (*FBH1*), **(C)** *RIBOSOMAL PROTEIN L27* (*RPL27*), **(D)** *JMJD5.2*, **(E)** *BASIC LEUCINE ZIPPER4* (*bZIP4*), and **(F)** *30S RIBOSOMAL PROTEIN S1* (*RPS1*) measured in the leaf +1 (L1, continuous green line), internodes 1 and 2 (I1, red dashed line), and internode 5 (I5, yellow dash-dotted line) of field-grown sugarcane by oligoarrays. Time series were normalized using *Z*-score. The light-gray boxes represent the night period.

**Supplemental Figure 13**

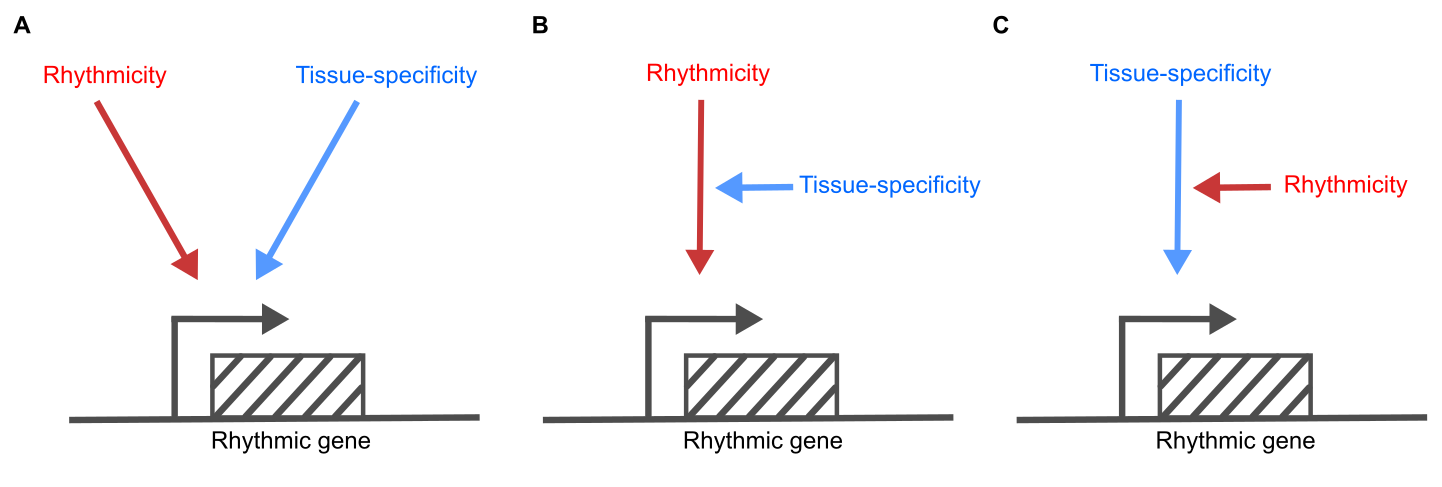

**Figure S13 – At least two different pathways are necessary to regulate tissue-specific rhythmic transcription.** One pathway confers rhythmicity to the transcript levels (red), while the other pathway confers tissue specificity (blue). These two pathways could regulate transcription independently **(A)**, or one could regulate the other **(B, C)**.

**Table S1 – Sugarcane genes and GenBank IDs**

| **Gene Symbol** | **Gene Name** | **SAS** | **GenBank ID** |
| --- | --- | --- | --- |
| *bZIP4* | *BASIC LEUCINE ZIPPER 4* | SCCCCL4005C09.g | CA094172.1 |
| *CM* | *CHORISMATE MUTASE* | SCCCCL4006E08.g | CA094273.1 |
| *CRY1.1* | *CRYPTOCHROME 1.1* | SCRUAD1133D10.b | CA217707.1 |
| *CRY2.1* | *CRYPTOCHROME 2.1* | SCRFST1042F05.g | CA178248.1 |
| *COMT* | *CATECHOL-O-METHYLTRANSFERASE* | SCEQLB2018G06.g | CA262225.1 |
| *CWI* | *CELL WALL INVERTASE* | SCVPRT3086D06.g | CA269857.1 |
| *ELF3* | *EARLY FLOWERING 3* | SCEZLB1009F09.g | CA113166.1 |
| *EXP* | *EXPANSIN* | SCCCCL5072C04.g | CA095540.1 |
| *FBA2* | *FRUCTOSE-BISPHOSPHATE ALDOLASE 2* | SCRULR1020D11.g | CA125940.1 |
| *FBH1* | *FLOWERING BASIC HELIX-LOOP-HELIX 1* | SCCCLR1022C05.g | CA119653.1 |
| *GAPDH* | *GLYCERALDEHYDE-3-PHOSPHATE DEHYDROGENASE* | SCQGAM2027G09.g | CA086777.1 |
| *GDH2* | *GLUTAMATE DEHYDROGENASE 2* | SCJLRT1021C04.g | CA135753.1 |
| *GI* | *GIGANTEA* | SCJFAD1014B07.b | CA067312.1 |
| *HB24* | *HOMEOBOX PROTEIN 24* | SCEQRT2095E01.g | CA139249.1 |
| *HDA6* | *HISTONE DEACETYLASE 6* | SCEPAM1024H12.g | CA073167.1 |
| *JMJD5.1* | *JUMONJI-C (JMJC) DOMAIN-CONTAINING PROTEIN 5.1* | SCEPRZ3086F09.g | CA156744.1 |
| *JMJD5.2* | *JUMONJI-C (JMJC) DOMAIN-CONTAINING PROTEIN 5.2* | SCCCLB1002G12.g | CA110901.1 |
| *LHY* | *LATE ELONGATED HYPOCOTYL* | SCCCLR1048E10.g | CA167119.1 |
| *MYB4* | *MYELOBLASTOSIS (MYB) DOMAIN PROTEIN 4* | SCCCLR1C05B03.g | CA189812.1 |
| *P4H* | *PROLYL 4-HYDROXYLASE* | SCCCST3006H07.g | CA180585.1 |
| *PAL* | *PHENYLALANINE AMMONIA LYASE* | SCEQRT1024E12.g | CA132523.1 |
| *PHYA.1* | *PHYTOCHROME A.1* | SCCCCL3080H06.g | CA093547.1 |
| *PHYB* | *PHYTOCHROME B* | SCQSLR1040D12.g | CA124822.1 |
| *PRR59* | *PSEUDO-RESPONSE REGULATOR 59* | SCACLR1057G02.g | CA116370.1 |
| *PRR73* | *PSEUDO-RESPONSE REGULATOR 73* | SCACLR1057C07.g | CA116387.1 |
| *PRR95* | *PSEUDO-RESPONSE REGULATOR 95* | SCCCLR1077F09.g | CA120437.1 |
| *RPL27* | *RIBOSOMAL PROTEIN L27* | SCAGLR2026E10.g | CA128063.1 |
| *RPS1* | *30S RIBOSOMAL PROTEIN S1* | SCBFST3135D12.g | CA181688.1 |
| *S15A* | *40S RIBOSOMAL PROTEIN S15* | SCJFRZ2007B06.g | CA151179.1 |
| *SMC1* | *STRUCTURAL MAINTENANCE OF CHROMOSOMES 1* | SCEZLB1008B03.g | CA113041.1 |
| *SPSII* | *SUCROSE-PHOSPHATE SYNTHASE II* | SCAGRT2037G07.g | CA137278.1 |
| *SuSy4* | *SUCROSE SYNTHASE 4* | SCEPCL6023F02.g | CA097247.1 |
| *SUT2* | *SUCROSE TRANSPORT PROTEIN 2* | SCEPLR1008A12.g | CA120749.1 |
| *SWEET11* | *SWEET11* | SCCCRT2002G08.g | CA137196.1 |
| *SWEET2* | *SWEET2* | SCSGRT2065C08.g | CA145445.1 |
| *TLP7* | *TUBBY-LIKE PROTEINS* | SCCCCL4001D08.g | CA093881.1 |
| *TOC1* | *TIME OF CAB EXPRESSION 1* | SCCCSB1002H04.g | CA167119.1 |
| *ZTL.1* | *ZEITLUPE* | SCCCLR1C07F05.g | CA190027.1 |

* Sugarcane Assembled Sequence.

**Table S2 – Sugarcane primers pairs used to validate oligo arrays expression levels using RT-qPCR**

| **Gene Symbol** | **SAS** | **Oligonucleotide sequence (5’ -> 3’)** | **Reference** |
| --- | --- | --- | --- |
| *FBA2* | SCRULR1020D11.g | FWD TGACACCATGGACCATGCAT  RVS GGCGACTAGCTCGATCTTTTCA | Rocha et al. (2007)^a^ |
| *HDA6* | SCEPAM1024H12.g | FWD GAGAAAGAGGAGGACATGGACAA  RVS CATAAGCTCCTCCGCTCCATA | Papini-Terzi et al. (2005)^b^ |
| *LHY* | SCCCLR1048E10.g | FWD CCACCACGGCCTAAAAGAAA  RVS TGGTTTTGTTGACTTGTCATTTGG | Hotta et al. (2011)^c^ |
| *MYB4* | SCCCLR1C05B03.g | FWD GGCAACAAATGGTCCCTGAT  RVS GCGTGTTCCAGTAGTTCTTGATCTC | Papini-Terzi et al. (2005)^b^ |
| *PAL* | SCEQRT1024E12.g | FWD CTTCCAGGGCACTCCCATT  RVS GAGAACTGCGCGAACATGAG | Papini-Terzi et al. (2005)^b^ |
| *PRR59* | SCACLR1057G02.g | FWD GACCCAGTTTTCCAACCCAAT  RVS CCCTCCGTGCTACTGTCCAA | Hotta et al. (2011)^c^ |
| *PRR73* | SCACLR1057C07.g | FWD CAAGTAATTCACCCCAAATCAGAGATA  RVS TCCCATAGATTCATCTTTATTCTCCTTAT | Hotta et al. (2011)^c^ |
| *PRR95* | SCCCLR1077F09.g | FWD CACCGATGGCATCCCTATTC  RVS TCTTGCCACATGGATGTTTTG | Hotta et al. (2011)^c^ |
| *TLP7* | SCCCCL4001D08.g | FWD GAGGCAGATGGCGAATGC  RVS GGACGATGCCTTTACAATGGA | Papini-Terzi et al. (2005)^b^ |
| *TOC1* | SCCCSB1002H04.g | FWD TTCTGCCTGAATTTGGCAAGTG  RVS GGCATCGAGCACACCAATGC | Hotta et al. (2011)^c^ |

^a^ Rocha et al. (2007) *BMC Genomics* 8:71, <https://doi.org/10.1186/1471-2164-8-71>

^b^ Papini-Terzi et al. (2005) *DNA Research* 12:27, <https://doi.org/10.1093/dnares/12.1.27>

^c^ Hotta et al. (2013) *PloS one* 8:e71847, https://doi.org/10.1371/journal.pone.0071847
